# C-LTMRs mediate wet dog shakes via the spinoparabrachial pathway

**DOI:** 10.1101/2024.06.10.597395

**Authors:** Dawei Zhang, Josef Turecek, Seungwon Choi, Michelle Delisle, Caroline L. Pamplona, Shan Meltzer, David D. Ginty

**Affiliations:** Department of Neurobiology, Howard Hughes Medical Institute, Harvard Medical School, 220 Longwood Avenue, Boston, MA 02115, USA

## Abstract

Mammals perform rapid oscillations of their body- “wet dog shakes” -to remove water and irritants from their back hairy skin. The somatosensory mechanisms underlying this stereotypical behavior are unknown. We report that Piezo2-dependent mechanosensation mediates wet dog shakes evoked by water or oil droplets applied to hairy skin of mice. Unmyelinated low-threshold mechanoreceptors (C-LTMRs) were strongly activated by oil droplets and their optogenetic activation elicited wet dog shakes. Ablation of C-LTMRs attenuated this behavior. Moreover, C-LTMRs synaptically couple to spinoparabrachial (SPB) neurons, and optogenetically inhibiting SPB neuron synapses and excitatory neurons in the parabrachial nucleus impaired both oil droplet- and C-LTMR-evoked wet dog shakes. Thus, a C-LTMR– spinoparabrachial pathway mediates wet dog shakes for rapid and effective removal of foreign particles from back hairy skin.

## Introduction

Wet dog shakes are an evolutionarily conserved behavior observed widely across mammalian species. This behavior consists of rapid oscillations of the head and upper trunk, typically following exposure of the back hairy skin of animals to water and other irritating or potentially damaging stimuli (*1-3*). Bouts of wet dog shakes can remove up to 70% of the water from a wet furry animal’s coat (*1*). Despite the prevalence of wet dog shakes and range of stimuli that can elicit them (*4-10*), the neurobiological mechanisms underlying this highly conserved, stereotypical behavior have remained elusive.

More than twelve physiologically and morphologically distinct primary somatosensory neuron subtypes innervate the hairy skin of mammals (*11*). These cutaneous sensory neurons collectively detect and encode a range of environmental stimuli, including mechanical, thermal, and chemical stimuli, with individual subtypes exhibiting unique stimulus-response profiles (*12-14*). Even simple mechanical stimuli, including static (step) indentation or brushing of the skin, can activate several mechanosensory neuron subtypes that exhibit distinct response properties. One noteworthy comparison is between Aβ RA-LTMRs, which form lanceolate endings that envelope the large guard and awl/auchene hair follicle types, and C-LTMRs, which form lanceolate endings around awl/auchene hairs as well as the most abundant hair follicle type, the zigzag hairs of the animal’s undercoat (*15*). Hairy skin Aβ RA-LTMRs are fast conducting, rapidly adapting mechanoreceptors that fire at the onset and offset of static indentation and respond strongly to rapid brushing of the skin, rendering them well-suited to function as dynamic touch detectors (*12, 15, 16*). In contrast, C-LTMRs are slow conducting, slowly to intermediately adapting mechanoreceptors that exhibit sustained responses to static indentation and brush, an after-discharge, and strong responses to slowly moving stimuli (*12, 15, 17, 18*). Therefore, C-LTMRs are poised to report the persistence of gentle, sustained mechanical stimuli acting on hairy skin. The contributions of these and other hair follicle-associated LTMRs and other cutaneous sensory neuron subtypes to the wet dog shake response is unknown.

How somatosensory signals encoding stimuli acting on mammalian furry skin are transformed in the central nervous system into motor commands that coordinate the stereotypical wet dog shake behavior is also an open question. All somatosensory neuron subtypes terminate within specific laminae of the spinal cord dorsal horn where they synapse onto different combinations of spinal cord interneurons and projection neurons. Spinal projection neurons then convey this somatosensory information to other CNS regions generate motor commands, autonomic responses, and sensory perception (*19, 20*). Prominent among dorsal horn projection neurons are the postsynaptic dorsal column neurons and the anterolateral tract projection neurons, which convey somatosensory signals encoding distinct features of the external world to different regions of the brain (*19, 20*). The identity of dorsal horn neurons and their corresponding output pathways that initiate wet dog shakes are unknown. Here, we have used mouse genetic, physiological, and behavioral approaches to uncover the somatosensory neurobiological basis of wet dog shakes.

## Results

### Oil droplets applied to the neck trigger wet dog shakes via a Piezo2-dependent mechanism

Wet dog shakes are observed widely across mammalian species in response to different stimuli (*4-10*). To begin to elucidate somatosensory mechanisms underlying this stereotypical behavior, we first tested a range of somatosensory stimuli for their ability to evoke wet dog shakes in mice. We observed that swimming in a water bath, spraying water onto the animal’s fur (*1, 3*) **(fig. S1A)**, applying a sunflower seed oil droplet, gentle air puff, or thin von Frey filaments to the back of the neck, and chloroquine injection into neck hairy skin all triggered wet dog shakes **(Fig. 1A, fig. S1B-D)**. Similar to the water bath and water spray stimuli, sunflower seed oil droplets applied to the back of the neck evoked wet dog shake bouts with comparable body motion kinetics in nearly all animals tested **(Fig. 1B, D, fig. S1E, F, movie S1)**. Because of its reliability, ease of application, and precise spatiotemporal control, we used the oil droplet stimulus to investigate the contribution of different somatosensory neuron subtypes and central circuitry underpinnings of this stereotypical mammalian behavior **(Fig. 1A)**.

**Fig. 1.**
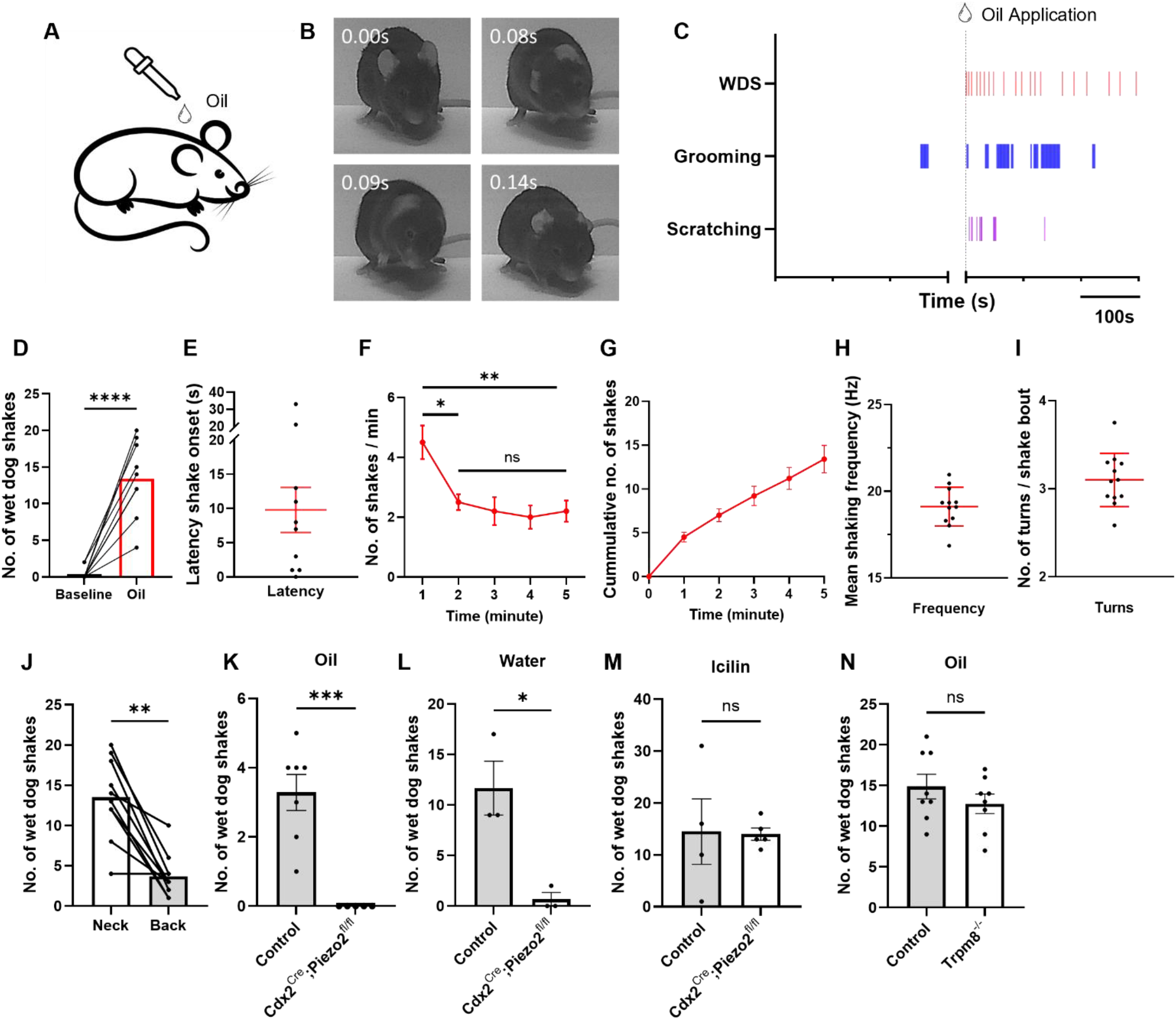
Oil droplets trigger wet dog shakes in mice via Piezo2-mediated mechanosensation. **(A)** Schematic diagram showing oil droplet application to the neck of a mouse. **(B)** Representative high-speed video frames of oil droplet-evoked wet dog shakes. Time is shown relative to wet dog shake onset. **(C)** Representative raster plots indicating episodes of wet dog shakes (WDS), grooming, and scratching behaviors before and after oil droplet application for one animal. Dashed line indicates oil droplet application time. Gap on the x-axis indicates a brief transition time between experiment sessions. **(D)** Total number of wet dog shakes before and after oil droplet application across animals (C57BL/6, N = 10) over 5 minutes. Comparisons were done using a two-sided paired t-test. **(E)** Wet dog shake onset latencies across animals shown in (D). **(F)** Average number of wet dog shakes over time following oil droplet treatment across animals shown in (D). Comparisons were done using one-way ANOVA across different time points. p-values adjusted for multiple testing using Tukey correction. **(G)** Cumulative plot of the average number of wet dog shakes across animals shown in (D) over time. **(H)** Average shaking frequencies and frequency variations across animals (N = 12). The oscillatory body movements are approximated as sine wave trajectories. Frequencies represent averages of at least 3 shaking bouts per animal. Error bar: SD. **(I)** Average number of oscillatory turns per shaking bout. The numbers of turns are averages of at least 3 shaking bouts per animal. These are the same shaking bouts presented in (H). **(J)** Total number of induced wet dog shakes when oil droplets were applied to the neck versus the back (N = 10) over 5 minutes. Comparison was done using two-sided paired t-test. **(K)** Total number of oil droplet-induced wet dog shakes in control (N = 7) and Piezo2 conditional knockout animals (N = 5) over 5 minutes. Control animals lack either the *Cre* allele or one copy of the floxed *Piezo2* allele. Comparison was done using two-sided unpaired t-test. **(L)** Total number of water-induced wet dog shakes in control (N = 3) and *Piezo2* conditional knockout animals (N = 3) over 5 minutes. Controls are of the same genotypes as in (K). Comparison was done using a two-sided unpaired t-test. **(M)** Total number of icilin-induced wet dog shakes in control (N = 4) and *Piezo2* conditional knockout animals (N = 5, same animals as in (K)) over 15 minutes. Comparison was done using a two-sided unpaired t-test. **(N)** Total number of oil droplet-induced wet dog shakes in control (C57BL/6, N = 8) and *Trpm8* knockout animals (N = 8) over 5 minutes. Comparison was done using a two-sided unpaired t-test. Shown are the means ± SEM, except in (H, I). ns p > 0.05, ^*^ p ≤ 0.05, ^**^ p < 0.01, ^***^ p < 0.001, ^****^ p < 0.0001. All black dots represent individual animals.

Oil droplets applied to the back of the neck induced many bouts of wet dog shakes over a five-minute recording period, and this was often accompanied by scratching and grooming **(Fig. 1C, D, fig. S1G, K)**. On average, a single oil droplet evoked wet dog shakes with a latency of ∼10 seconds, although this varied across animals and trials from less than one second to over half a minute **(Fig. 1E)**. Animals responded most robustly during the first minute following an oil droplet application and subsequently performed fewer shakes **(Fig. 1F, G)**. In contrast, grooming behaviors persisted throughout the five-minute recording window, whereas the timing of scratching varied across individuals **(fig. S1H, I, L, M)**. Individual wet dog shake episodes were highly stereotyped across animals, with a mean frequency of oscillatory movements of 19.12 Hz (SD = 1.12) and an average of 3.10 (SD = 0.31) full back and forth turns per shaking bout **(Fig. 1H, I)**. Animals were more responsive to oil droplets applied to the back of the neck, displaying significantly more wet dog shakes and scratching bouts, compared to the same stimulus applied to the lower back **(Fig. 1J, fig. S1J, N)**.

Stimulus-evoked wet dog shakes may reflect responses to different sensory modalities (*4-10, 21*). Indeed, wet dog shakes evoked by liquids including water or oil droplets may reflect responses to mechanical or thermal stimuli or both (*3*). To test whether mechanosensation is the primary driver of water and oil droplet-induced wet dog shakes (*3*), the mechanosensitive ion channel Piezo2 was deleted in all DRG neurons below the upper cervical region of the body using *Cdx2*^*Cre*^*;Piezo2*^*fl/fl*^ animals (*16, 22, 23*), and responses to water or oil droplet application to the neck were examined in mutants and controls. We observed an absence of oil droplet-evoked wet dog shakes **(Fig. 1K)** and a nearly complete loss of water-evoked wet dog shakes in the *Piezo2* mutants **(Fig. 1L)**. Importantly, the lack of responses in *Piezo2* mutant mice is due to insensitivity to mechanical stimuli rather than an inability to perform wet dog shakes because intraperitoneal injection of icilin, which is an agonist of the cold/cool-sensitive ion channel Trpm8 (*24-26*), evoked wet dog shakes to a comparable extent in *Piezo2* mutants and control mice **(Fig. 1M, fig. S1O, P)**. In contrast, animals lacking Trpm8 (*Trpm8*^*-/-*^) (*27*) and wildtype controls exhibited comparable bouts of wet dog shakes evoked by oil droplets **(Fig. 1N)**, whereas icilin-induced wet dog shakes were lost in *Trpm8*^*-/-*^ animals (*28*) **(fig. S1Q)**. Thus, water- and oil droplet-evoked wet dog shakes require Piezo2-dependent mechanosensation.

### Low threshold mechanoreceptors are preferentially activated by oil droplets applied to hairy skin

We next sought to identify the primary mechanosensory neurons that mediate mechanically evoked wet dog shakes induced by oil droplets. To achieve this, a series of *in vivo* DRG calcium imaging experiments in isoflurane-anesthetized mice was conducted to assess responses across the principal mechanosensory neuron subtypes following application of oil droplets to hairy skin **(Fig. 2A)**. Due to the physical challenges of accessing cervical DRGs that innervate hairy skin of the neck and unavoidable damage to the skin and sensory nerve endings during cervical DRG surgical exposure, we conducted experiments on lumbar L4 DRGs and applied oil droplets to hairy skin of the thigh. We examined the responsiveness of six distinct DRG mechanosensory neuron types, including four low threshold mechanoreceptors (LTMRs): Aβ RA-LTMRs, Aβ SA1-LTMRs, Aδ-LTMRs, and C-LTMRs; along with the two most mechanically sensitive C-fiber high threshold mechanoreceptors (HTMRs): C-HTMRs expressing MrgprB4 (MrgprB4^+^) and C-HTMRs expressing MrgprD (MrgprD^+^) **(*13*)**. In agreement with previous findings (*12, 13, 16, 29-32*), all four LTMR types and both C-HTMR types responded to brushing or pinching of the skin **(fig. S2)**. Application of oil droplets to hairy skin evoked robust responses in most C-LTMRs, Aδ-LTMRs, and Aβ SA1-LTMRs **(Fig. 2B-D)**, while few Aβ RA-LTMRs, C-HTMRs (MrgprB4^+^), and C-HTMRs (MrgprD^+^) responded to the stimulus **(Fig. 2E-G)**. Averaging population responses across the sensory neuron types showed that C-LTMRs, Aδ-LTMRs, and Aβ SA1-LTMRs exhibited robust responses with distinct temporal dynamics **(Fig. 2H, I)**, whereas Aβ RA-LTMRs, C-HTMRs (MrgprB4^+^), and C-HTMRs (MrgprD^+^) exhibited virtually no responses to the oil droplet stimulus **(Fig. 2H)**. Thus, C-LTMRs, Aδ-LTMRs, and Aβ SA1-LTMRs are candidate drivers of mechanical stimulus-evoked wet dog shakes.

**Fig. 2.**
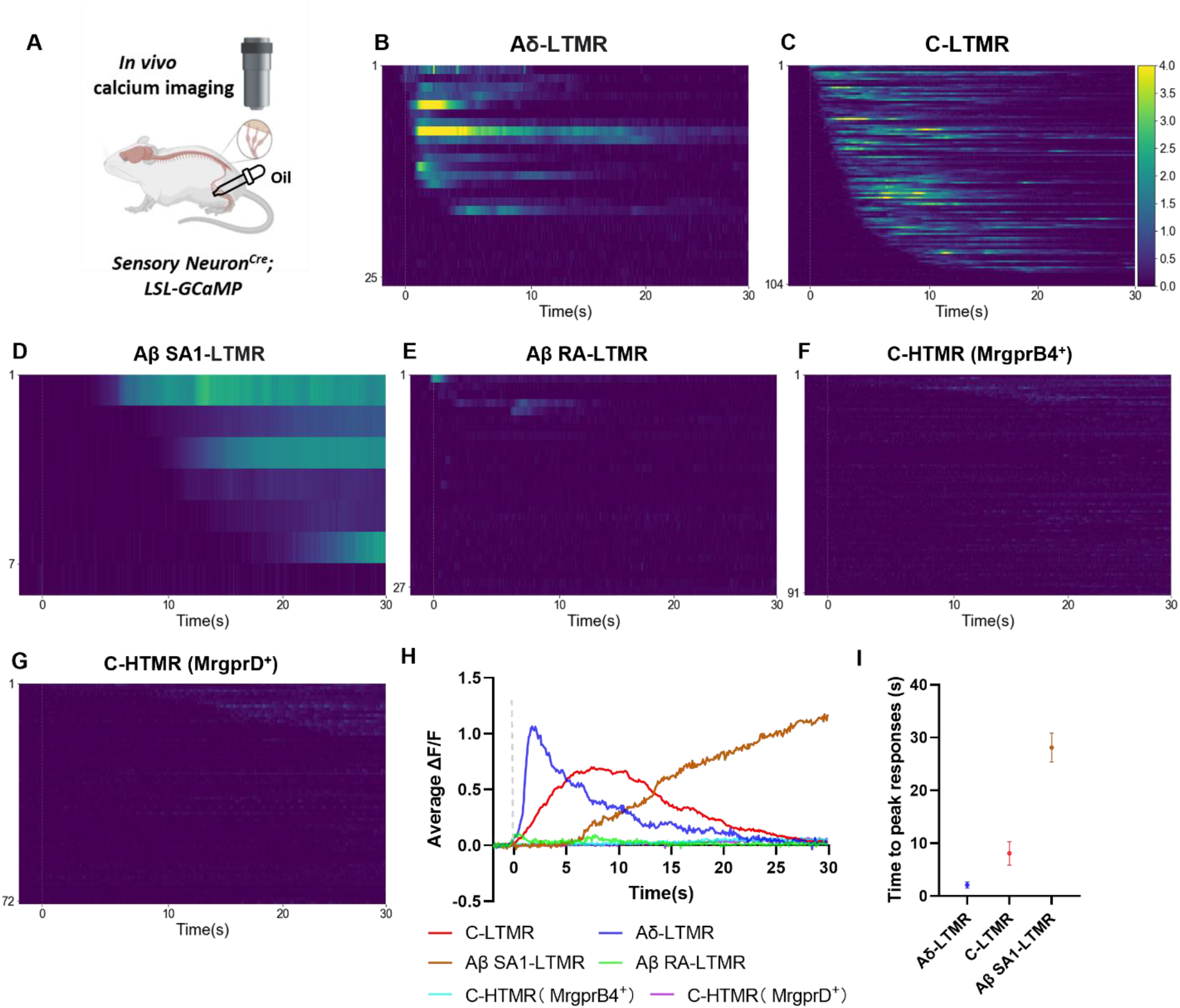
Select LTMR subtypes are preferentially activated by oil droplets applied to hairy skin. **(A)** Schematic diagram of *in vivo* DRG calcium imaging experiments with oil droplet stimuli. **(B-G)** Heatmaps of calcium responses in (B) Aδ-LTMRs (*TrkB*^*CreER*^; *Ai96*, N = 3, n = 25), (C) C-LTMRs (*TH*^*CreER*^; *Ai148*, N = 3, n = 104), (D) Aβ SA1-LTMRs (*TrkC*^*CreER*^; *Ai148*, N = 2, n = 7), (E) Aβ RA-LTMRs (*Ret*^*CreER*^; *Ai148*, N = 3, n = 27), (F) C-HTMRs (MrgprB4^+^), (*MrgprB4*^*CreER*^; *Ai148*, N = 3, n = 91), and (G) C-HTMRs (MrgprD^+^) (*MrgprD*^*CreER*^; *Ai148*, N = 3, n = 72) in response to oil droplet. Rows of the heatmap represent responses of individual DRG neurons over time. **(H)** Average calcium responses of Aδ-LTMRs, C-LTMRs, Aβ SA1-LTMRs, Aβ RA-LTMRs, C-HTMRs (MrgprB4^+^), and C-HTMRs (MrgprD^+^) in response to oil droplets applied to the skin. **(I)** Time to reach peak calcium responses in Aδ-LTMRs, C-LTMRs, and Aβ SA1-LTMRs after oil droplet application. Shown are the means ± SEM. Dashed lines indicate the oil droplet application time. N: number of animals. n: number of cells.

### Optogenetic stimulation of C-LTMRs induces wet dog shakes

We next sought to determine whether stimulation of one or more LTMR subtypes is sufficient to evoke wet dog shakes. Therefore, we expressed the light-activated cation channel ReaChR (*33, 34*) in the four LTMR subtypes and activated them by illuminating skin of the neck or back while monitoring mice for the wet dog shake behavior **(Fig. 3A, B)**. Remarkably, optogenetic stimulation of neurons labeled using *TH*^*T2a-CreER*^, which are C-LTMRs and sympathetic neurons, consistently elicited wet dog shakes under both continuous and 10 Hz light stimulation paradigms **(Fig. 3C, movie S2)**. These responses were mediated by C-LTMRs and not sympathetic neurons because optogenetic stimulation of sympathetic neurons alone, using a Flpo driver line (*DBH*^*p2a-Flpo*^) that labels sympathetic neurons but not C-LTMRs, failed to evoke wet dog shakes **(fig. S3)**. Surprisingly, optogenetic stimulation of other LTMR subtypes, including Aδ-LTMRs, Aβ SA1-LTMRs, and Aβ RA-LTMRs was insufficient to elicit wet dog shakes **(Fig. 3C)**, despite our findings that these optical stimuli can evoke robust neural responses in the spinal cord (*35, 36*).

**Fig. 3.**
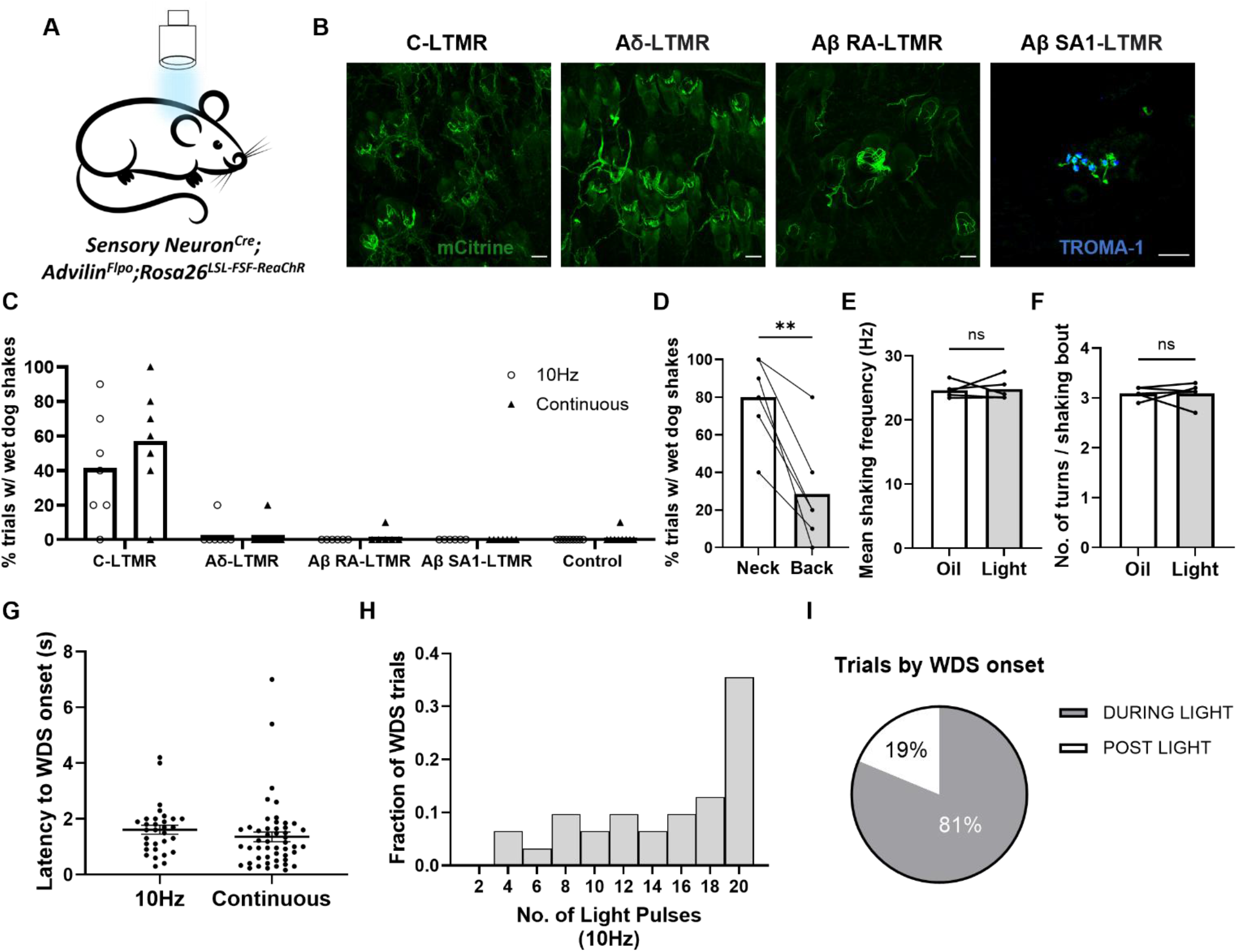
Optogenetic stimulation of C-LTMRs, but not other LTMRs, is sufficient to trigger wet dog shakes. **(A)** Schematic diagram showing optogenetic stimulation of primary sensory neuron types innervating the skin using blue laser light. **(B)** Representative images showing immunostaining-enhanced ReaChR expression (labeled by mCitrine) in the cutaneous axonal endings of C-LTMRs (labeled by *TH*^*CreER*^; *Adv*^*Flpo*^; *Rosa26*^*LSL*- *FSF-ReaChR*^), Aδ-LTMRs (labeled by *TrkB*^*CreER*^; *Adv*^*Flpo*^; *Rosa26*^*LSL-FSF-ReaChR*^), Aβ RA-LTMRs (labeled by *Ret*^*CreER*^; *Adv*^*Flpo*^; *Rosa26*^*LSL-FSF-ReaChR*^), and Aβ SA1-LTMRs (labeled by *TrkC*^*CreER*^; *Adv*^*Flpo*^; *Rosa26*^*LSL-FSF-ReaChR*^). Scale bars, 50 μm. **(C)** Percentage of trials with wet dog shakes induced by optogenetic activation of different LTMR types across animals. Same labeling strategies as in (B). Number of animals tested for each genotype: *TH*^*CreER*^; *Adv*^*Flpo*^; *Rosa26*^*LSL-FSF-ReaChR*^, N = 7. *TrkB*^*CreER*^; *Adv*^*Flpo*^; *Rosa26*^*LSL-FSF-*^ _*ReaChR*_, N = 6. *Ret*^*CreER*^; *Adv*^*Flpo*^; *Rosa26*^*LSL-FSF-ReaChR+*^, N = 6. *TrkC*^*CreER*^; *Adv*^*Flpo*^; *Rosa26* ^*LSL-FSF-ReaChR*^, N = 7. Controls are littermates without CreER or Flpo allele ( N = 9). **(D)** Percentage of trials with wet dog shakes induced by continuous light exposure (2s) on the neck versus back in *TH*^*CreER*^; *Adv*^*Flpo*^; *Rosa26*^*LSL* -*FSF-ReaChR*^ animals (N = 6, including 2 ReaChR allele homozygotes). Comparison using two-sided paired t test. **(E)** Average shaking frequency of oil- and light-induced wet dog shakes in *TH*^*CreER*^; *Adv*^*Flpo*^; *Rosa26*^*LSL-FSF-ReaChR*^ animals (N =5). Frequencies were averaged from at least 3 shaking bouts for each animal. Comparison using two-sided paired t test. **(F)** Average number of oscillatory turns per shaking bout induced by oil and light stimuli across animals. The number of turns were averaged from at least 3 shaking bouts for each animal. Same shaking bouts as in (E). Comparison using two-sided paired t test. **(G)** Latency to wet dog shake (WDS) initiation after light onset using 2 seconds of 10Hz pulse light or continuous light stimuli. Number of trials for 10Hz and continuous light are 31 and 49, respectively. Dots represent individual trials. Shown are the means ± SEM. **(H)** Histogram of the number of pulses (10Hz) applied prior to the onset of induced wet dog shakes. Only trials with pulse light-induced wet dog shakes were included in the quantification. Number of trials = 31. **(I)** Percentage of trials in which wet dog shakes were initiated during and after the light stimuli, respectively. Only trials with successful light induction of wet dog shakes were included in the quantification. Number of trials = 80. Bars represent means. All markers represent individual animals except in (G). ns p > 0.05, ^**^ p < 0.01.

We further explored C-LTMR-evoked wet dog shake responses and found that, as with oil droplet-evoked responses, optogenetic stimulation of C-LTMRs terminating in dorsal neck hairy skin triggered more pronounced wet dog shakes compared to stimulation of C-LTMRs terminating in hairy skin of the lower back **(Fig. 3D)**. This difference in behavioral responses was not due to variations in C-LTMR density, terminal morphology, or genetic labeling specificity or efficiency across the two skin regions, as the innervation density and gross central termination patterns of C-LTMRs innervating neck and lower back hairy skin were indistinguishable **(fig. S4)**. Moreover, characteristic features of wet dog shakes, including shaking frequency and the number of body turns per bout, were comparable between those evoked by oil droplet application and C-LTMR photostimulation **(Fig. 3E, F)**. Additionally, as with oil droplet evoked responses, the onset latency of wet dog shakes following C-LTMR optostimulation varied across trials and ranged from less than half a second to seven seconds **(Fig. 3G)**. Interestingly, wet dog shakes were most frequently initiated towards the end of the 10 Hz light stimulation sessions **(Fig. 3H)** and in some cases commenced only after cessation of light stimulation (19% of trials) **(Fig. 3I)**. This delayed onset is notably longer than at least some other reflex behaviors induced by C-fiber photoactivation, which can prompt paw withdrawal within approximately 700 milliseconds (*37*). The delayed onset may imply involvement of C-LTMR information processing in supraspinal regions of the nervous system for initiation of wet dog shakes.

### Ablation of C-LTMR neurons reduces wet dog shakes

Because of the robust behavioral responses observed following C-LTMR optogenetic stimulation, we next asked whether C-LTMRs are necessary for wet dog shakes induced by the oil droplet stimulus. To this end, we generated a new mouse line *Tafa4*^*CreER*^ **(fig. S5A)**, which labels C-LTMRs but not sympathetic neurons (*38*) **(fig. S5B-D)**, and crossed this line to *R26*^*eGFP*- *DTA*^ mice (hereafter referred to as TAFA4-DTA mice) to ablate C-LTMRs. Five consecutive days of tamoxifen treatment of adult TAFA4-DTA mice led to ∼92% reduction of C-LTMRs, as determined by TH antibody staining of DRGs. Moreover, C-LTMRs were selectively lost using this ablation strategy, as DRG neuronal populations marked by IB4^+^, CGRP^+^, and NFH^+^ did not differ from control mice lacking the *Tafa4*^*CreER*^ allele **(Fig. 4A, B)**.

**Fig. 4.**
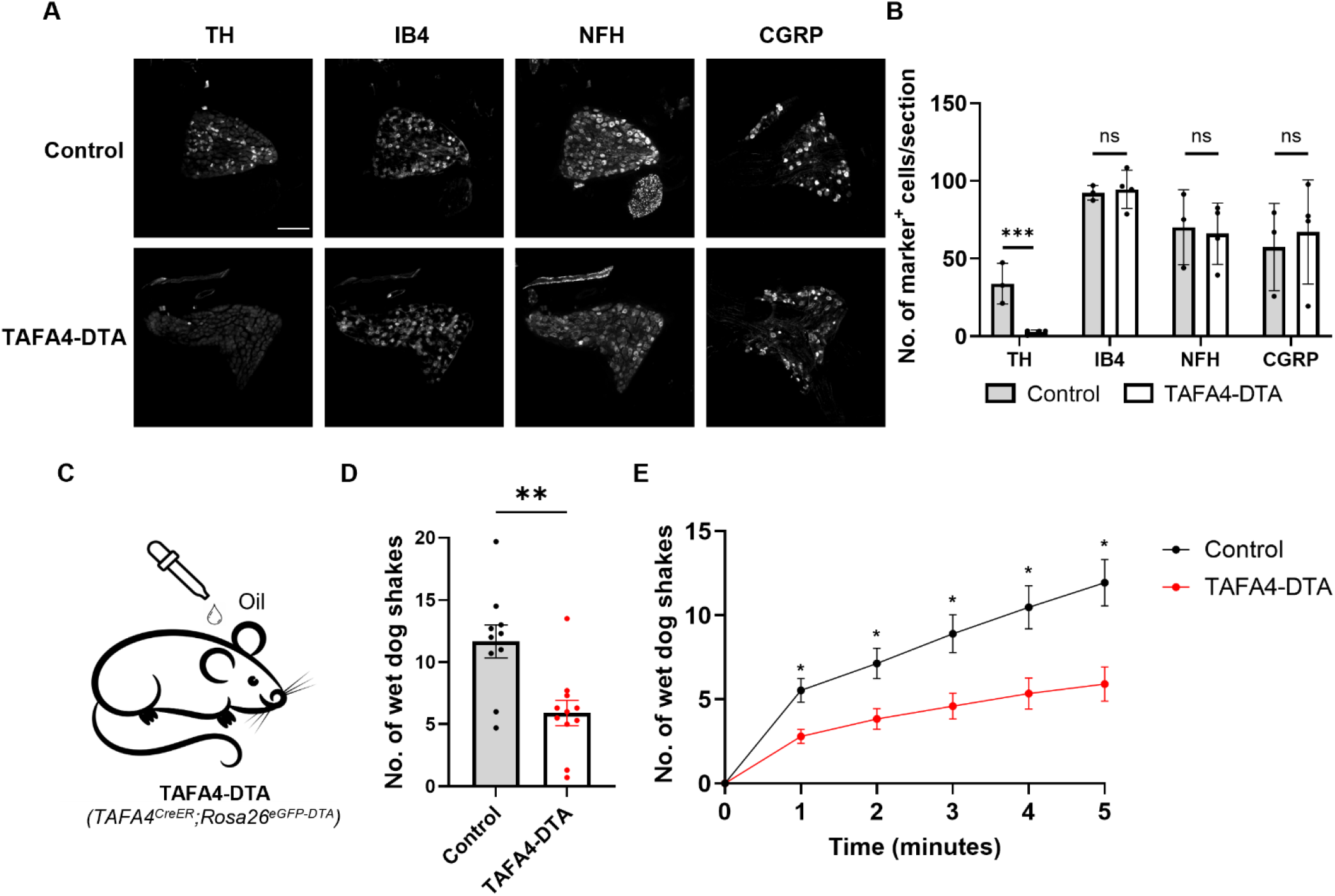
Ablation of C-LTMRs reduces oil droplet-evoked wet dog shakes. **(A)** Representative images of DRG immunostaining of control (*Rosa26*^*LSL-DTA*^) and TAFA4-DTA (*Tafa4*^*CreER*^; *Rosa26*^*LSL-DTA*^) animals. Scale bars, 200 μm. **(B)** Quantification of the number of cervical DRG neurons stained with cell type markers exemplified in (A). The number of cells per section was averaged from at least 5 sections for each animal. Each dot represents an animal. Control, N = 3; TAFA4-DTA, N = 4. Comparison using two-sided unpaired t-test. **(C)** Schematic diagram showing oil droplet application to the neck of control and TAFA4-DTA animals. **(D)** Average number of wet dog shakes induced by oil droplet application in littermate control (N = 10) and TAFA4-DTA animals (N = 11). Control animals lack the *Tafa4*^*CreER*^ allele. Each dot represents the averaged response from at least 2 repeated trials of an animal. Comparison using two-sided unpaired t-test. **(E)** Cumulative plot of the number of wet dog shakes over time. Comparison using two-way ANOVA. p-values adjusted for multiple testing using Bonferroni correction. Shown are the means ± SEM. ns p > 0.05, ^*^ p≤0.05, ^**^p < 0.01, ^***^p < 0.001.

We observed a significant reduction (∼50%) in oil droplet-evoked wet dog shakes in TAFA4-DTA mice compared to controls, a finding consistent across trials **(Fig. 4C-E, fig. S7A)**. To mitigate potential confounds associated with TAFA4 expression elsewhere in the body **(fig. S5F)**, we also ablated C-LTMRs along with other C-fiber neurons and some Aδ-fiber HTMRs in the peripheral nervous system using *Scn10a*^*Cre*^ *;R26*^*eGFP-DTA*^ animals (hereafter referred to as SNS-DTA mice) (*39*) and tested their responses to oil droplet stimulation **(fig. S6A, B)**. SNS-DTA mice mirrored the TAFA4-DTA animals, exhibiting a comparable reduction (∼48%) in oil-induced wet-dog shakes **(fig. S6C, D)**, suggesting that the reduction of responses seen in TAFA4-DTA animals was due to the loss of C-LTMRs.

To further assess somatosensory and motor behaviors in TAFA4-DTA mice, we performed a series of additional behavioral measurements. Our experiments revealed normal locomotor activity in these animals using the open field and balance beam assays **(fig. S7B, C)**. Moreover, icilin-induced wet-dog shakes were normal in TAFA4-DTA mice **(fig. S7D)**, indicating that the wet dog shake motor pattern generation function remains intact in these mice. Also, TAFA4-DTA mice did not exhibit differences in grooming behaviors in response to oil droplets compared to their littermate controls **(fig. S7E)**. Notably, TAFA4-DTA mice exhibited a reduction (∼58%) in oil-induced scratching behaviors compared to controls **(fig. S7F)**, but these mice did not exhibit alterations in scratching in response to histamine or chloroquine **(fig. S7G-H)**. In addition, TAFA4-DTA mice exhibited normal cold thermosensation, which was assessed using a temperature preference assay **(fig. S7I)**, and these mutants also showed normal paw withdrawal responses to von Frey filaments applied to their hind paw **(fig. S7J, K)**. Collectively, these findings demonstrate that C-LTMRs contribute to oil droplet-evoked wet dog shakes but are not required for a range of other somatosensory behaviors. Residual wet dog shake responses observed in TAFA4-DTA mice **(Fig. 4C-E, fig. S7A)** may reflect sufficiency of the small number (∼8%) of remaining C-LTMRs in these mutants or the contribution of one or more other LTMRs in evoking the behavior.

### The spinoparabrachial pathway mediates wet dog shakes

The finding that C-LTMRs contribute to stimulus-evoked wet dog shakes afforded an opportunity to ask how these enigmatic mechanosensory neurons engage central circuits to mediate somatosensory behaviors. C-LTMRs are known to terminate in lamina IIiv of the superficial spinal cord dorsal horn (*15, 17*). Therefore, we asked whether C-LTMR signals are conveyed from the superficial dorsal horn to the lateral parabrachial nucleus (PBN_L_) to mediate wet dog shakes. To investigate this possibility, we first asked whether optogenetic activation of the central terminals of C-LTMRs could evoke postsynaptic responses in spinoparabrachial (SPB) neurons. We used whole-cell patch-clamp electrophysiological recordings of NK1R^+^ and GRP83^+^ SPB neurons (SPNs), which together account for approximately 90% of superficial SPB neurons, in spinal cord slices (*40*) **(Fig. 5A)**. We found that C-LTMRs are indeed synaptically coupled to SPNs, since optogenetic activation of C-LTMR terminals evoked excitatory postsynaptic currents (EPSCs) in both SPN populations **(Fig. 5B, C)**.

**Fig. 5.**
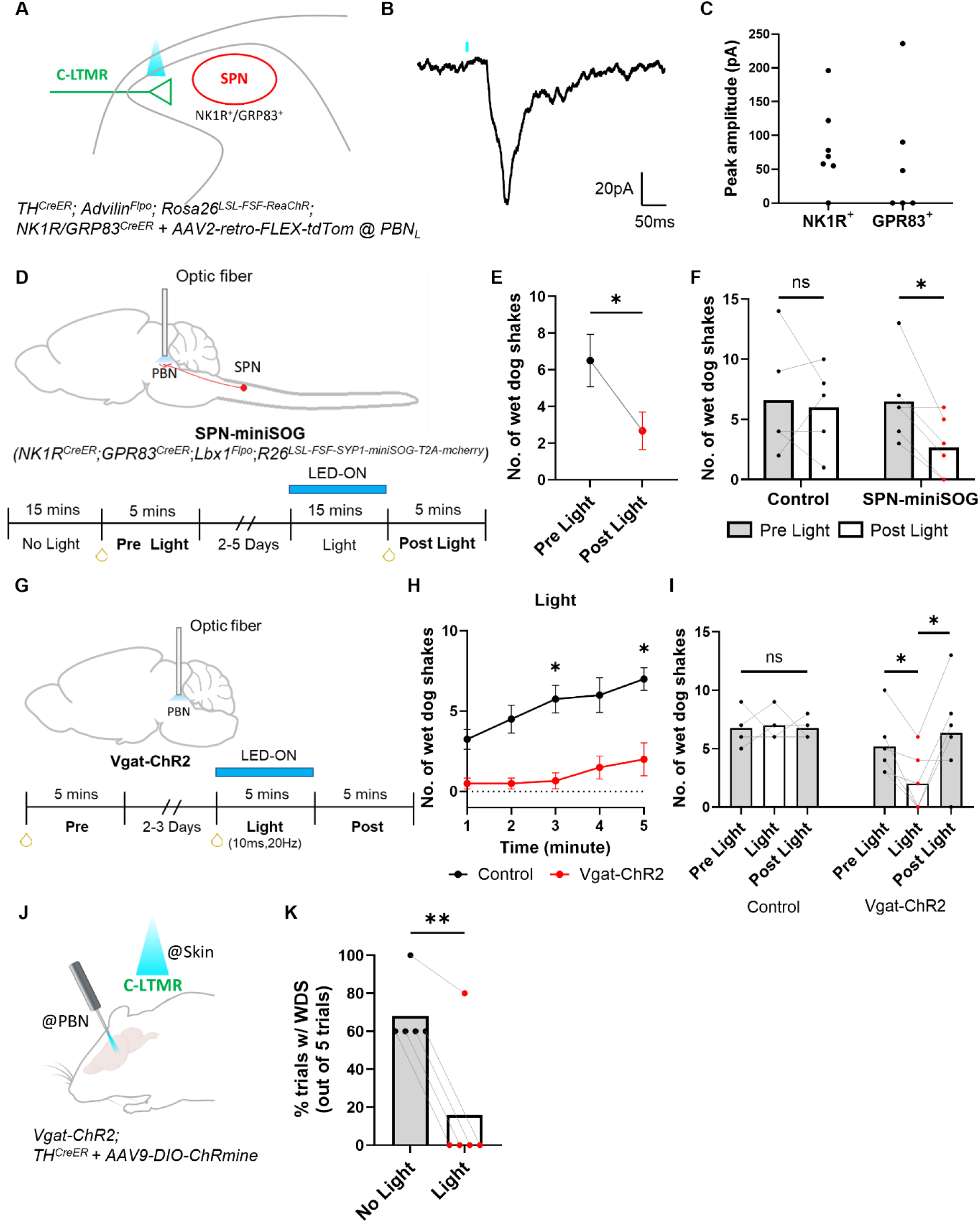
The spinoparabrachial pathway mediates oil droplet- and C-LTMR-evoked wet dog shakes. **(A)** Schematic diagram showing the genetic labeling strategy and coronal spinal cord slice electrophysiological recordings from SPB neurons (SPNs) during blue light stimulation of TH^+^ primary afferent terminals. **(B)** Representative trace of a light activated EPSC from an SPN. The blue line indicates the light onset. **(C)** Peak amplitudes of light induced EPSCs in NK1R^+^ and GPR83^+^ SPNs. Dots represent individual cells (NK1R^+^, n = 8; GPR83^+^, n = 7). **(D)** Schematic diagram showing optogenetic inhibition of synaptic release from SPNs and the experimental design. **(E)** Average number of oil-induced wet dog shakes before and after SPN terminal light stimulation in SPN-miniSOG animals (N = 6) over a 5 minute measurement period. Shown are the means ± SEM. **(F)** Total number of oil-induced wet dog shakes before and after light stimulation in littermate control (N = 5) and SPN-miniSOG animals over a 5 minute measurement period. Control animals lack either the *Lbx1*^*Flpo*^ allele or the *R26*^*LSL-FSF-SYP1-miniSOG-mcherry*^ allele. The same wet dog shake bout measurements as in (E) for SPN-miniSOG animals. **(G)** Schematic diagram showing optogenetic inhibition of PBN neurons using Vgat-ChR2 animals and the experimental design. **(H)** Cumulative plot of the average number of wet dog shakes in control (C57Bl/6, N = 4) and Vgat-ChR2 animals (N = 6) during PBN light stimulation. Total light power used was 3 mW (power density 23.86 mW/mm2). Comparisons were done using two-way ANOVA. p-values adjusted for multiple testing using Bonferroni correction. **(I)** Total number of oil-induced wet dog shakes before, during, and after light stimulation in control and Vgat-ChR2 animals from (H). Dots represent individual animals. Bars represent means. Comparisons were done using a two-sided paired t-test. **(J)** Schematic diagram showing the genetic labeling strategy and experimental design for simultaneous blue light activation of C-LTMR endings in the skin and Vgat^+^ inhibitory neurons on the PBN. **(K)** Average percentage of trials with light (on the skin) induced wet dog shakes (WDS) with and without (‘Light’ vs. ‘No Light’) Vgat-ChR2 photoactivation in the PBN across animals (N = 5). N: number of animals. n: number of cells. The same SPN-miniSOG animals were used in (E-H). All dots represent individual animals except in (C) and (E). Bars represent means. Comparisons were done using a two-sided paired t-test. ^*^ p ≤ 0.05.

To ask whether SPNs mediate mechanically evoked wet dog shake behaviors, we used the light-activated protein miniSOG (*41*) to inhibit SPN synaptic transmission in the PBN_L_. Fusion of miniSOG to synaptophysin (SYP1) enables light-triggered oxidation of synaptic proteins necessary for triggering neurotransmitter release and synaptic transmission (*41*). We generated a dual recombinase-dependent mouse line, *R26*^*LSL-FSF-SYP1-miniSOG*^ that enables expression of SYP1-miniSOG at synaptic terminals **(fig. S8A)** and thus inhibition of synaptic transmission using optical stimuli applied to axon terminals. Consistent with previous findings in worms (*41*), light application to spinal cord excitatory neuron axon terminals expressing SYP1-miniSOG suppressed excitatory drive by approximately 75% **(fig. S8B-D)**. Therefore, we expressed SYP1-miniSOG in SPNs using *NK1R*^*CreER*^; *Gpr83*^*CreER*^; *Lbx1*^*Flpo*^; *R26*^*LSL-FSF-SYP1-miniSOG*^ mice (hereafter referred to as SPN-miniSOG mice) **(*40*)** and implanted optical fibers into the PBN to enable light-evoked inhibition of SPN synaptic transmission in awake, freely moving animals **(Fig. 5D)**. Upon light stimulation of the PBN, SPN-miniSOG mice exhibited a reduction (∼58%) in oil droplet evoked wet dog shakes **(Fig. 5E, F)**. In contrast, oil droplet evoked wet dog shakes were unperturbed following light stimulation of the PBN in control animals lacking SYP1-miniSOG expression. Moreover, SPN-miniSOG mice exposed to light exhibited a trend towards a reduction in oil droplet-induced scratching behaviors (∼75%, p=0.06) and a less pronounced trend in grooming behaviors, consistent with prior findings that the SPB projection pathway mediates itch sensation (*42, 43*) **(fig. S9A-D)**. Light stimulation of SPN-miniSOG animals did not alter back hairy skin tactile sensitivity, measured using a tactile pre-pulse inhibition (tPPI) assay (*44, 45, 57*) **(fig. S9E, F)**.

To complement the findings using SPN-miniSOG mice and determine whether C-LTMR mediated wet dog shakes require the SPB pathway, we silenced the PBN directly, through optogenetic stimulation of GABAergic neurons of the PBN using Vgat-ChR2 animals and assessed the consequences of this manipulation on oil droplet and C-LTMR evoked wet dog shakes. Consistent with the SPN-miniSOG manipulations, broad inhibition of the PBN through optogenetic stimulation of PBN local and projecting GABAergic neurons inhibited wet dog shakes evoked by both oil droplets **(Fig. 5G-I)** and photostimulation of C-LTMR terminals in the skin **(Fig. 5J, K, fig. S9G)**. Collectively, these findings demonstrate the contribution of a C-LTMR–spinoparabrachial pathway in mechanically evoked wet dog shakes.

## Discussion

Wet dog shakes are an evolutionary conserved behavior observed across mammals that serves to remove water and other irritants from back and neck hairy skin, a skin region that is largely unreachable by self-grooming or -licking. Here, we report that a C-LTMR–spinoparabrachial pathway mediates mechanically-evoked wet dog shakes.

The contributions of C-LTMRs to perception and behavior has been a matter of considerable interest. A prevailing view is that human C-mechanoreceptors mediate affective, pleasurable touch, an idea stemming from human patient studies and correlational analyses of neuronal activity and reports of pleasantness (*46-48*). C-LTMRs, which in mice form lanceolate endings around vellus hair follicles and terminate in spinal cord lamina IIiv (*15*), are the most sensitive of the C-fiber mechanoreceptor subtypes and are often considered the murine equivalent of the C-mechanoreceptors described in humans (*13, 48, 49*). However, in mice other C-fiber mechanoreceptor types with higher force thresholds and that do not innervate hair follicles may also contribute to affective touch (*30, 50*). Although C-LTMRs have an inferred role in affective touch, our findings implicate them as mediators of wet dog shakes. It is noteworthy that in the seminal work identifying C-LTMRs, nearly 80 years ago (*51*), Zotterman suggested that these C-fiber mechanoreceptors may underlie the sensation of ‘tickle.’ Zotterman suggested that the ultrasensitive C-LTMR responses to gentle forces acting on the skin and their after-discharge properties align with the observation that in humans the lightest detectable forces evoke a tickle percept, which may persist following cessation of the stimulus (*51*). In support of Zotterman’s notion, we observed a reduction in both oil droplet evoked wet dog shakes and scratching in animals lacking C-LTMRs. It is also noteworthy that C-LTMRs form lanceolate axonal endings exclusively around the small vellus hairs of the body (*15*), which comprise the undercoat, an insulating layer of hair that may serve to protect against environmental threats including water, insects, and other parasites. Related to this, recent findings have implicated C-LTMRs in promoting tissue repair (*52*). Thus, we propose that C-LTMRs detect the lightest forces acting on hairy skin, including water, movements of insects and parasites, and other stimuli that deflect vellus hairs, triggering motor behaviors that have evolved to remove water, mechanical irritants, and potential threats. Interestingly, despite a uniform distribution of C-LTMR terminals in neck and lower back hairy skin, the biased and more robust response to oil droplet and C-LTMR photostimulation of dorsal neck hairy skin suggests the existence of unique motor pattern generating circuitry associated with the cervical spinal cord, which warrants future investigation.

Prior work reported that electrical stimulation of the hippocampus and septum can induce wet dog shakes (*53*). However, the involvement of peripheral and ascending somatosensory pathways that initiate this stereotypical motor behavior had remained unknown. Our findings show that SPB neurons play a key role in conveying C-LTMR signals to the brain to initiate wet dog shakes. Notably, photoactivation of at least two other C-fiber sensory neurons with distinct response properties, the C-HTMR (MrgprB4^+^) and C-Cold (Trpm8^+^) neurons, can also evoke wet dog shakes **(fig. S10)**. This is consistent with the observation that wet dog shakes can be elicited by stimuli that span different sensory modalities, including cooling agents and pruritogens **(fig. S1D, Q)**. Because the SPB pathway is implicated in both thermosensation and itch (*40, 42, 43, 54, 55*), we suspect that the PBN mediates wet dog shakes evoked by different sensory modalities. Future research will aim to reveal the PBN neuronal ensembles that drive wet dog shakes, and how PBN signals are relayed to downstream brain regions and central motor pattern generation centers to initiate this conserved, stereotypical behavior.

## Acknowledgements

We thank Celine Santiago and Jia Yin Xiao for providing *Piezo2* conditional knockout mice, members of the Ginty lab for discussions, and Kitwa Ng, Lijun Qi, Rosa Martinez-Garcia, Andrew Shuster, Zoe Sarafis, Celine Santiago, and Anda Chirila for comments on the manuscript. We thank Caiying Guo and the Gene Targeting and Transgenics Facility at Janelia Research Campus for generating the *Tafa4*^*CreER*^ and *R26*^*LSL-FSF-SYP1-miniSOG-mcherry*^ mouse lines. We thank the Boston Children’s Hospital Viral Core for packaging AAV viruses.

## Author contributions

DZ and DDG conceived the study. DZ performed and analyzed behavioral, calcium imaging, histology, and optogenetic manipulation experiments. JT and SC performed and analyzed the slice physiology experiments and helped with the generation and characterization of the *R26*^*LSL-FSF-SYP1-miniSOG-mcherry*^ mouse line. MD performed tPPI experiments and assisted with other behavioral experiments. CLP helped with the analyses of behavioral experiments and supported genotyping. SM contributed to the generation and initial characterization of the *Tafa4*^*CreER*^ mouse line. DZ and DDG wrote the paper with input from all authors.

## Data and materials availability

All data are available in the manuscript or the supplementary material. Reagents and code are available from the corresponding author upon reasonable request.

## Supplementary Materials

Materials and Methods

Figs. S1 to S10

Movies S1 to S2

