## Supplementary Materials for "C-LTMRs mediate wet dog shakes via the spinoparabrachial pathway"

### Materials and Methods

#### Animals

All experimental procedures in this study were approved by the Harvard Medical School Institutional Animal Care and Use Committee (IACUC) and were performed in compliance with the Guide for Animal Care and Use of Laboratory Animals. Both males and females were used in this study. The animals used in this study that were published previously includes C57BL/6J (JAX #664), CD1 (Charles River #22), $Ret^{CreER}$ (MGI:4437245), $TrkB^{CreER}$ (MGI:5616440),$TrkC^{CreER}$ (MGI:5707203), $TH^{CreER}$ (MGI:5569743), $MrgprD^{CreER}$ (MGI:5924939), $MrgprB4^{Cre}$ (MGI:5448562), $Cdx2^{Cre}$ (JAX #009350), $Scn10a^{Cre}$ (MGI:3053096), $Nk1R^{CreER}$ (MGI:7539708), $GPR83^{CreER}$ (MGI:7539709), $Advilin^{Flpo}$ (MGI:6740440), $Lbx1^{Flpo}$ (MGI:5770780), $DBH^{p2a\text{-}Flpo}$ (MGI:5770774), Ai14 (JAX #007914), Ai96 (JAX #028866), Ai148 (JAX #030328), $Rosa26^{LSL\text{-}ReaChR\text{-}mCitrine}$ (JAX #026294), $Rosa26^{LSL\text{-}FSF\text{-}ReaChR\text{-}mCitrine}$ (JAX # 024846), $Rosa26^{FSF\text{-}ReaChR\text{-}mCitrine}$ (derived from JAX #024846), VGAT-ChR2-EYFP (JAX # 014548), $Rosa26^{eGFP\text{-}DTA}$ (JAX #032087), $Piezo2^{fl/fl}$ (JAX #027720), and Trpm8 KO (JAX #008198). Newly published animals in this study include $Tafa4^{T2a\text{-}CreER}$ and $Rosa26^{FSF\text{-}LSL\text{-}SYP1\text{-}miniSOG\text{-}mcherry}$. These mice were generated at the Janelia Research Campus Gene Targeting and Transgenic Facility using CRISPR-based homologous recombination techniques in embryonic stem (ES) cells. Chimeras were produced via blastocyst injection, and germline transmission was confirmed through standard tissue genotyping PCR. All animals used for experiments carrying reporter alleles or VGAT-ChR2-EYFP were heterozygous for these alleles unless mentioned otherwise.

#### Tamoxifen treatment

Tamoxifen (Sigma T5648) was first dissolved in 100% ethanol to a concentration of 20 mg/mL and then mixed at a 1:1 ratio with sunflower seed oil (Sigma S5007). In cases where tamoxifen was delivered embryonically, progesterone (Sigma P0130) was added to the ethanol along with tamoxifen at a 1:2 ratio. All tamoxifen mixtures were vortexed and then vacuum centrifuged to evaporate the ethanol.

| Genotype | Time | Dose, frequency |
|---|---|---|
| $Ret^{CreER}$; (Flpo); Reporter | E10.5-E11.5 | 3mg, once |
| $TrkC^{CreER}$; (Flpo); Reporter | E12.5 | 3mg, once |
| $TrkB^{CreER}$; (Flpo); Reporter | P21-P28 or P5 | 2mg or 0.5mg, once |
| $TH^{CreER}$; (Flpo); Reporter | P21-P28 | 2mg, once |
| $MrgprD^{CreER}$; Reporter | >P13 | 2mg, once |
| $Tafa4^{CreER}$; Ai14 | >4 weeks | 2mg, once |
| $Tafa4^{CreER}$; $Rosa26^{eGFP\text{-}DTA}$ | >4 weeks | 2mg, 5 consecutive days |
| $Nk1R^{CreER}$; $GPR83^{CreER}$; $Lbx1^{Flpo}$; $Rosa26^{FSF\text{-}LSL\text{-}SYP1\text{-}miniSOG\text{-}mcherry}$ | P21-P28 | ~4mg (0.2mg/g), once |

#### AAV Viral Delivery

All AAV viruses were produced and packaged at the Boston Children's Hospital Viral Core Facility. For systemic AAV delivery, 10 µl of AAV9-Ef1a-DIO-ChRmine-mScarlet-WPRE (5.99E+14 vg/mL) (Addgene #130998) viruses were injected intraperitoneally into neonatal (P0-P3) mice or intrathecally into adult mice (aged over 6 weeks). After neonatal viral delivery, tamoxifen was administered for 3-5 consecutive days at weaning age. Following adult viral delivery, two doses of tamoxifen (2 mg/animal) were administered intraperitoneally, one day and three days post-injection. For AAV injections into the brain, mice (aged 6–10 weeks) were secured in a stereotaxic frame (Kopf Instruments) and anaesthetized with continuous inhalation of isoflurane (1.5–2.5%) during the surgery. Between 100 and 300 nl of AAV2-retro-CAG-FLEX-tdTomato-WPRE (2.22E+13 gc/ml) viruses were injected into the PBN using pulled glass pipettes (Drummond Wiretrol II) and a microsyringe pump injector (World Precision Instruments UMP3). Two or three doses of tamoxifen were administered via oral gavage 5–14 days after the virus injections.

**Behavioral Assays**

*Induction and Measurement of Wet Dog Shakes*

Animals were habituated in black matte acrylic chambers (H* W * L = 12 cm x 12 cm x 12 cm) for at least 15 minutes before any stimulation. For oil droplet stimuli, 20-25 µl of sunflower seed oil (Sigma S5007) were applied to either the neck or the back of the animals using a glass Pasteur pipette (VWR# 14673-010). For most experiments, the oil droplet was applied to the predefined skin region while the animals were partially restrained, with their tails held by the experimenter, after which the animals were immediately returned to the chamber for recording. For experiments involving the quantification of behavior onset latency and comparison of baseline and stimulated behavior activities, the oil droplet was applied while the animals were resting in the chambers without restraint. For water stimuli, each animal was placed in a cage filled with warm water set to a depth that allowed the animals' feet to barely touch the ground as they floated. Water was also gently splashed onto the dorsal side of the animals by hand. Afterwards, the animals were transferred to the acrylic chambers for behavioral recordings.

For chemical stimuli, icilin (Sigma #I9532) was suspended in 1% Tween 80/saline and injected intraperitoneally (1 mg/ml, 100 µL); chloroquine (200 µg in 50 µL saline, Sigma #C6628) and histamine (100 µg in 30 µL, Sigma #H7125) were injected intradermally/subcutaneously at the nape of the neck using 31g BD insulin syringes. For both the air puff and von Frey stimuli, the hairs on animals' necks were shaved prior to habituation. Different pressures of air puff and varying forces of von Frey hairs were then applied to the animals' necks. All types of stimuli were administered to mice aged over 3 weeks. The behaviors of each animal were video recorded following stimulus application at 30fps, 240fps or 300fps. Behavioral events, including wet dog shakes, scratches, and grooming, were manually annotated and quantified using the recorded videos.

*Open Field*

Animals were habituated in black matte acrylic open field chambers (W * L = 25 cm x 25 cm) for two days. On the testing day, each animal was placed in the chamber and allowed to explore freely for 5 minutes, with the session being recorded on video. The travel distance and movement trace maps for each animal were analyzed and generated using customized MATLAB code.

*Balance Beam*

The balance beam was constructed from a 1-meter matte white acrylic bar with a flat 12 mm surface, mounted 50 cm above the benchtop. Black matte acrylic boxes were placed at both ends of the beam to serve as the start and end enclosures for the animals. A camera mounted above the balance beam captured the animals' crossings. Prior to the test, animals were accustomed to the setup and trained to cross the beam for two days until they could do so freely. If animals refused to cross, the investigator would gently touch their hindquarters to encourage forward movement. Animals were tested to cross the beam four times on the test day. The time taken to cross the beam was manually quantified based on the recorded video.

*Temperature Preference*

Animals were group habituated for 5 minutes on a metal plate in a black matte acrylic chamber (30 cm x 15 cm) divided by an arched doorway in the middle. The metal plate was placed on a heating pad set to 30°C during habituation. Habituation occurred over two days before testing. The test setup consisted of two cold plate coolers (TE Tech #CP-121HT) placed side by side; the gap between the plates was filled with a customized acrylic bar. The temperature of each plate was controlled by two independent temperature controllers (TE Tech #TC-720). The black matte acrylic chamber was positioned on top of the two plates, with the doorway at the center. A camera was installed on the ceiling for video capture. On test day 1, both plates were set to 30°C. Each animal had 5 minutes to explore the chamber. On test day 2, the temperatures were set to 30°C and 18°C, allowing animals to explore for another 5 minutes. The time animals spent on each side of the chamber/each plate was analyzed using customized MATLAB code.

*Von Frey Test*

Mice were placed in clear plastic chambers on an elevated wire mesh and habituated for at least 1 hour. The plantar surface of the hind paw was stimulated with a set of calibrated von Frey monofilaments. To determine the percentage of paw withdrawal responses, stimulation was applied five times with the same force, and responses were converted to a percentage. The up-down assay (*56*) was used to determine the paw withdrawal threshold..

*Tactile Pre-pulse Inhibition*

tPPI was performed as described previously using the San Diego Instruments startle reflex system (SR-LAB Startle Response System) (*44, 45, 57*). Briefly, a tactile prepulse (0.9-psi air puff, 50 ms) was delivered at variable interstimulus intervals (50, 100, 250, 500, and 1000 ms) before the acoustic startle pulse. The mouse’s startle response was measured using an accelerometer. Following a 5-minute acclimation phase, testing sessions included a) acoustic pulse trials alone, b) air puff alone, and c) pseudorandomly mixed trials of prepulse/pulse, pulse alone, and no stimulation. Maximal accelerometer responses (mV) were recorded in a 100 ms window following the end of each stimulus. Percent PPI was calculated as %PPI = [1 – (startle response/pulse alone response)] × 100. The response to the air puff alone was measured as (prepulse alone response/pulse alone response) – (no stimulation response/pulse alone response).

### ***In vivo* Calcium Imaging**

Mice (aged 3-8 weeks) were anesthetized with 1-3% inhalational isoflurane on a customized acrylic platform during surgeries and imaging sessions. Their body temperature was monitored and maintained at 37°C using a temperature controller (Warner Instruments #TC-344B) and a thermoelectric heater (Honeywell #C3200-6145). Incisions were made over the L4 and L5 vertebral spines and nearby paravertebral muscles to allow a customized aluminum spinal clamp to secure the spine. The overlaying muscles and bones of the L4 spine were then removed using spring scissors and forceps. Surgifoam sponges (McKesson #1972) and Q-tips® cotton swabs were used to control bleeding. DRGs were imaged using a 470 nm LED under an epifluorescence microscope (Zeiss Axio Examiner) with a 10X air objective (Zeiss Epiplan, NA=0.20). Images were captured with a CMOS camera (Thorlabs # CS505MU1) at a frame rate of 10 Hz or 20 Hz.

All stimuli were applied to the hairy skin near the thigh region in the following order: brush, oil droplet, and pinch. Brush stimulation was applied using a paintbrush (JOINREY size 2), oil droplets (25 µl) were applied using a glass Pasteur pipette (VWR #14673-010), and pinch stimulation was applied using metal forceps with dull tips (FST Dumont #2). Each type of stimulus was applied at multiple spots within the same skin region to cover the receptive fields for most cells in the imaging field of view. Stimuli were manually administered at a set time point after the baseline window.

Imaging analyses were performed using ImageJ and Python. All images were converted into 8-bit stacks, and then motion corrections were conducted using the moco plugin (*58*) (https://github.com/NTCColumbia/moco). Background subtraction was performed using ImageJ built-in function after motion correction to facilitate manual selection of the ROI. The analysis of ImageJ-extracted ROI (cell) fluorescent intensities was conducted using Python. Fluorescent intensities of cells were converted into ΔF/F, where F was defined as the average intensity during the baseline window (2 seconds before the set stimulation time). For each cell, the session displaying the highest peak ΔF/F value across multiple skin spots under the same type of stimulus was considered to represent the optimal response and was selected for further analysis. Only cells showing a calcium response to pinch or sustained static indentation (for SA1-LTMRs) were included in the analyses of brush and oil responses.

#### ***In vivo* Optogenetic Stimulation and Fiber-optic Cannula Implantation**

For hairy skin stimulation, mice over 6 weeks of age had the hairs on their neck and back shaved prior to testing. Mice were habituated in behavior chambers (H* W * L = 12 cm x 12 cm x 12 cm) for at least 15 minutes before light stimulation. A blue laser light (473nm) was manually directed onto the neck or the back of the resting animals (without active locomotion and behaviors such as grooming, scratching, or rearing) through a 1000-micron fiber patch cable (Thorlabs #M59L) attached to a collimator (Thorlabs #F240SMA-532). The light was either pulsed at 10 Hz (20 ms pulse width) or provided a consistent illumination for 2 seconds per trial. The illuminated area on the skin ranged between 15-30 mm in diameter, with a power density between 0.18-0.74 $mW/mm^2$. Each animal underwent 10 trials using the same stimulation paradigm. Animal behaviors were simultaneously captured by a front camera and an overhead camera at either 30 fps, 240 fps or 300fps. For C-HMTR ($MrgprD^+$) hindpaw stimulation, animals were briefly habituated in clear plastic chambers on elevated wire mesh for at least 15 minutes. The plantar surface of the hind paw was stimulated 5 times with the same light source (power density: 1.91 $mW/mm^2$) and stimulation paradigm described above.

For brain stimulation, optic fibers (400 micron in diameter, 0.66 NA, Doric #B280-4655-3.5) were bilaterally implanted into the PBN (from –5.2 to –5.0 mm posterior to bregma, ± 1.4 to 1.6 mm from the midline, and –2.5 to −2.7 mm ventral to the dura), following a previously described protocol (*40*). Mice aged 6–12 weeks at the time of implantation were given at least one week to recover before any behavioral tests. Prior to experiments, mice were briefly anesthetized with inhalational isoflurane to facilitate the attachment of branching fiber optic patch cords and zirconium sleeves (both from Doric Lenses) to implanted cannulas. Light stimulations were controlled by a programmable LED driver (Doric Lenses) through Doric Neuroscience Studio software. For miniSOG experiments, axon terminals in the PBN were stimulated with continuous blue LED (Doric #CLED_465, 465-470nm, total power of 3.5mW, power density of 27.87 $mW/mm^2$) for 15 minutes. For VGAT-ChR2 related experiments, inhibitory axon terminals in the PBN region were stimulated with a blue LED (Doric #CLED_465, 465-470nm, 20Hz, 10ms pulse width) with a total power of 3 mW (power density of 23.86 $mW/mm^2$) or 5 mW (power density of 39.77 $mW/mm^2$) during 'light' trials.

***In vitro* Slice Electrophysiology**

*ReaChR related experiments*

Similar to previously described (*40*), mice (5–7 weeks of age) were anesthetized before lumbar enlargements were dissected out in an ice-cold choline solution (in mM: 92 choline chloride, 2.5 KCl, 1.2 $NaH_2PO_4$, 30 $NaHCO_3$, 20 HEPES, 25 glucose, 5 sodium ascorbate, 2 thiourea, 3 sodium pyruvate, 10 $MgSO_4{\bullet}7H_2O$, 0.5 $CaCl_2{\bullet}2H_2O$) and mounted in 0.3% LMP agarose (Life Technology #16520-100). The lumbar spinal cords were sliced in the transverse plane (350 μm) (Leica VT1200S) and recovered at 34 °C for 30 min in oxygenated (95% $O_2$ and 5% $CO_2$) HEPES holding solution (in mM: 86 NaCl, 2.5 KCl, 1.2 $NaH_2PO_4$, 35 $NaHCO_3$, 20 HEPES, 25 glucose, 5 sodium ascorbate, 2 thiourea, 3 sodium pyruvate, 1 $MgSO_4{\bullet}7H_2O$, 2 $CaCl_2{\bullet}2H_2O$). After recovery, spinal cord slices were placed at RT for 30 min before recordings. Spinal cord slices were then superfused with oxygenated (95% $O_2$ and 5% $CO_2$) recording ACSF (in mM: 2.5 $CaCl_2{\bullet}2H_2O$, 1.0 $NaH_2PO_4{\bullet}H_2O$, 119 NaCl, 2.5 KCl, 1.3 $MgSO_4{\bullet}7H_2O$, 26 $NaHCO_3$, 25 glucose, 1.3 sodium L-ascorbate) at RT in a recording chamber mounted on a SliceScope Pro 6000 electrophysiology rig (Scientifica). Labelled SPNs within superficial lamina were identified using tdTomato fluorescence and patched under 40x objective using infrared differential interference contrast microscopy (Hamamatsu Photonics ORCA-Flash 4.0; Scientifica SliceScope Pro 6000). The pipette resistance ranged from 3 to 4 MΩ, and the electrodes were filled with an intracellular solution (in mM: 135 K-gluconate, 5 KCl, 0.5 $CaCl_2$, 5 EGTA, 5 HEPES, 5 MgATP, pH 7.2). Signals were acquired using a Multiclamp 700B amplifier (Molecular Devices). The data were low-pass filtered at 2 kHz, digitized at 10 kHz with an A/D converter (Molecular Devices Digidata 1440A) and stored using a data-acquisition program (Molecular Devices, Clampex version 10). The liquid junction potential was not corrected. Primary afferent synaptic terminals were stimulated with wide-field blue LED illumination through the 40× objective (CoolLED pE-300, 1 ms pulse width, light intensity = 27 mW). Cell capacitance, current amplitude, latency and jitter were analyzed using Clampfit (Molecular Devices, version 10).

*miniSOG related experiments*

Adult mice (>P30) of both sexes expressing miniSOG in excitatory neurons ($Vglut2^{IRES\text{-}Cre}$; $Cdx2^{Flp}$; $Rosa26^{LSL\text{-}FSF\text{-}SYP1\text{-}mini\text{-}SOG\text{-}mcherry}$), or no miniSOG ($Vglut2^{IRES\text{-}Cre}$; $Cdx2^{Flp}$) were anesthetized with isoflurane and transcardially perfused using room temperature choline chloride dissection solution of the following contents (in mM): 110 choline-Cl, 2.5 KCl, 1.25 $NaH_2PO_4$, 25 $NaHCO_3$, 25 glucose, 0.5 $CaCl_2$, 7 $MgCl_2$ , 3.1 Na pyruvate, 11.6 Na ascorbate, 0.002 (R)-CPP, 0.005 NBQX, oxygenated with 95% $O_2$ /5% $CO_2$. The spinal cord was removed in the dark, embedded in 4% low melting point agarose and 200 µm sagittal sections were made using a Leica 1200S vibratome. Slices were transferred to a light-shielded holding chamber with solution containing the following (in mM): 127 NaCl, 2.5 KCl, 1.25 $NaH_2PO_4$, 25 $NaHCO_3$, 25 glucose, 1.5 $CaCl_2$, 1 $MgCl_2$, and allowed to recover at 35 C for at least 30 minutes before being brought to room temperature.

Experiments were performed in the dark at room temperature in ACSF of the same composition as the holding solution described above. Whole-cell voltage-clamp recordings were made from random spinal cord dorsal horn (Laminae II-IV) neurons. Borosillicate electrodes (1.5-2 MΩ) for recording EPSCs were filled with the following solution (in mM): 130 Cs-Gluconate, 10 HEPES, 1 EGTA, 0.1 $CaCl_2$, 5 TEA, 1 QX-314, 0.2 D-600, pH 7.3 using CsOH. For recording IPSCs, pipettes were filled with solution (in mM): 110 CsCl, 10 HEPES, 10 EGTA, 10 Cs-BAPTA, 4 $CaCl_2$, 1 $MgCl_2$, 10 TEA, 2 QX-314, 0.2 D-600, pH 7.3 with CsOH. Recordings of EPSCs were performed in the presence of 2 µM (R)-CPP to block NMDARs. Recordings of IPSCs were performed in the presence of 5 µM NBQX and 2 µM (R)-CPP to block AMPARs and NMDARs, respectively. All recordings were performed at -60 mV, with liquid junction potential left unsubtracted. Series resistance (5-20 MΩ) was compensated up to 80%, with whole-cell capacitance compensated at 5 pF for all experiments.

To evoke EPSCs or IPSCs, a glass pipette of similar resistance to recording pipettes was placed on the same dorso-ventral level as the recorded cell at least 500 µm away (rostral or caudal) from the recorded cell body. Single electrical stimuli were delivered every 5 seconds, with the intensity set to deliver a short latency (2-4 ms) 200-500 pA EPSC or IPSC. Stimuli were evoked for 5 minutes to obtain baseline values. After 5 minutes, continuous blue light was delivered through a 40x objective directly over the cell body (10 $mW/mm^2$, measured at the objective) for 5-7 minutes. Light was generated by a cold white LED (Thorlabs MCWHL8-C1) and 470-20 nm band pass excitation filter.

For analysis, traces from the baseline period were averaged, and the evoked EPSC or IPSC was integrated to obtain the total charge. Traces from the post-stimulus period (2-10 minutes following the offset of light) were also averaged and integrated to obtain the total charge. Summarized values are represented as the total charge normalized to the baseline charge for each experiment.

**Immunohistochemistry**

The following primary antibodies and lectins were prepared: rabbit anti-TH (Sigma #AB152, 1:500), sheep anti-TH (Sigma #AB1542, 1:500), chicken anti-GFP (Aves Labs #GFP-1020, 1:1000), goat anti-GFP (US Biological Life Sciences #G8965-01E, 1:1000), goat anti-mCherry (CedarLane #AB0040-200, 1:1000), chicken anti-NFH (Aves #NFH, 1:500), rabbit anti-NF200 (Sigma #N4142, 1:500), mouse anti-PKCγ (Santa Cruz #sc-166385, 1:500), rat anti-Troma (DSHB TROMA-I supernatant, 1:100), rabbit anti-CGRP (Immunostar #24112, 1:500), and Isolectin B4 (Alexa 647 conjugated, Invitrogen). Secondary antibodies were Alexa 488, 546, or 647 conjugated and sourced from donkey or goat anti-rabbit, chicken, goat, rat, or mouse (Invitrogen or Jackson), prepared at a 1:500 dilution.

For DRG, spinal cord and brain staining, isoflurane anesthetized animals were transcranially perfused with PBS followed by 4% PFA. Tissues were dissected, post-fixed in 4% PFA overnight at 4°C, then washed with PBS. Spinal cord and brain tissues were cryoprotected in 30% sucrose overnight at 4°C, while DRG was treated for 1 hour at room temperature. Tissues were embedded in OCT (Fisher #14-373-65), frozen, sectioned into 25–40 micron slices using a cryostat, and mounted on Superfrost Plus slides (Fisher #12-550-15) for staining. After washing three times in 1x PBS and 0.1% PBST, and blocking for 1 hour at room temperature, tissues were incubated with primary antibodies overnight at 4°C. Secondary antibodies were applied for 1-2 hours at room temperature. Following three PBS washes, slides were mounted using Fluoromount-G (Southern

Biotech #0100-20 or 0100-01) and imaged with a fluorescent or confocal microscope (Zeiss LSM 700, Olympus VS120 slide scanners or Zeiss LSM 900).

For whole mount skin staining, animals were euthanized using $CO_2$ or perfused similarly to other tissues. Hair was removed with Nair™, and underlying fat tissues were scraped off. Skin samples were washed with PBS, fixed in 2% paraformaldehyde in PBS or Zamboni's fixative at 4°C for 1-2 hours, then cut into smaller pieces. Samples underwent multiple washes in 1x PBS and 1% PBST, before primary antibodies were applied in blocking solution (5% serum, 75% PBST, 20% DMSO) and incubated with gentle shaking at room temperature for 2-5 nights. After three times of washing in 1% PBST, tissues were incubated with secondary antibodies for 1-2 nights at room temperature. Following final washes, tissues were dehydrated in a methanol series (50%, 75%, 100%), cleared in BABB (1 part Benzyl Alcohol: 2 parts Benzyl Benzoate), and mounted on slides in BABB for confocal imaging.

#### ***In situ*** **Hybridization (RNA scope)**

Similar as previously described (*11*), we followed the ADCBio Manual RNAscope Assay user guide for sample preparation and treatment. Briefly, fresh frozen DRGs were used for cryosectioning, and 4% PFA was used for section fixation. Sections were digested with Protease IV for 15 minutes at room temperature before a series of probe incubations. Slides were mounted using Fluoromount-G (Southern Biotech #0100-01) and imaged using a fluorescent microscope (Zeiss LSM 700). Probes used in this study include Mm-Trpm8-C2 (ADC Bio #420451-C2).

#### **Statistical Analyses**

Statistical analyses were performed using GraphPad Prism. Data are presented as either mean ± SEM or mean ± SD, as indicated in the figure legends. The specific methods used for comparisons between two groups are detailed in the figure legends. Additionally, p-values for multiple comparisons were adjusted as described in the figure legends.

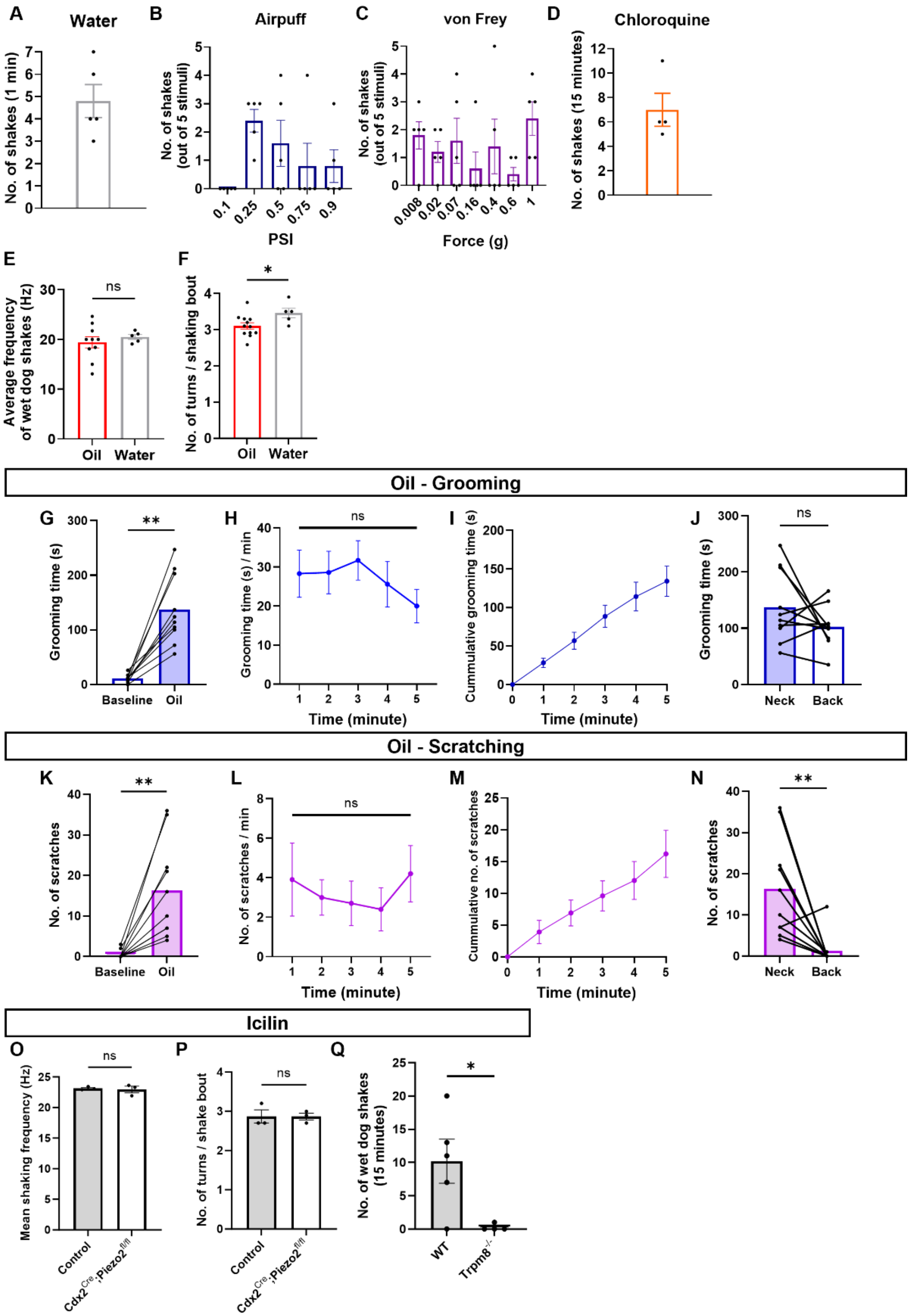


**Fig. S1. Wet dog shakes, grooming, and scratching induced by a range of sensory stimuli**
**(A)** Total number of wet dog shakes induced by water stimuli across C57BL/6 mice (N = 5) over 1 minute.

**(B)** Number of trials with air puff-induced wet dog shakes across C57BL/6 mice (N = 5) presented with air puffs using different air pressures. All animals had their neck hair shaved prior to the stimulation.
**(C)** Number of trials with von Frey filament-induced wet dog shakes across C57BL/6 mice (N = 5) stimulated with different filament forces. All animals had their neck hair shaved prior to the stimulation.
**(D)** Total number of chloroquine-induced wet dog shakes across C57BL/6 mice (C57BL/6, N = 4) over 15 minutes.
**(E)** Average shaking frequencies induced by oil (N = 10) and water (N = 5). Frequencies were averaged from at least 3 shaking bouts per animal. Comparison was done using a two-sided unpaired t-test.
**(F)** Average number of oscillatory turns per shaking bout induced by oil and water. The numbers of turns are averaged from at least 3 shaking bouts per animal. Comparison was done using a two-sided unpaired t-test.
**(G)** Total grooming amount before and after oil droplet application across animals (N = 10) over 5 minutes. Comparison was done using a two-sided paired t-test.
**(H)** Average grooming amount over time following oil droplet treatment across animals in (G). Comparison using one-way ANOVA across different time points. p-values adjusted for multiple testing using Tukey correction.
**(I)** Cumulative plot of the grooming amount across animals in (G) over time.
**(J)** Total grooming amount when oil droplets were applied to the neck versus the back (N = 10) over 5 minutes. Comparison was done using a two-sided paired t-test.
**(K)** Total number of scratching events before and after oil droplet application across animals (N = 10) over 5 minutes. Comparison was done using a two-sided paired t-test.
**(L)** Average number of scratching events over time following oil droplet treatment across animals in (K). Comparison using one-way ANOVA across different time points. p-values adjusted for multiple testing using Tukey correction.
**(M)** Cumulative plot of the number of scratching events across animals in (K) over time.
**(N)** Total number of scratching events when oil droplets were applied to the neck versus the back (N = 10) over 5 minutes. Comparison was done using a two-sided paired t-test.
**(O)** Average shaking frequencies induced by oil droplets in control (N = 3) and Piezo2 conditional knockout animals (N = 3). Control animals lack either the *Cre* allele or one copy of the floxed *Piezo2* allele. Frequencies were averaged from at least 3 shaking bouts per animal. Comparison was done using a two-sided unpaired t-test.
**(P)** Average number of oscillatory turns per shaking bout induced by oil. The numbers of turns were averaged from at least 3 shaking bouts per animal. Comparison was done using a two-sided unpaired t-test. These are the same shaking bouts presented in (O).
**(Q)** Total number of icilin-induced wet dog shakes in wildtype (C57BL/6, N = 5) and Trpm8 knockout animals (N = 4) over 15 minutes. Comparison was done using a two-sided unpaired t-test.
Shown are the means ± SEM. * $p \leq 0.05$, ** $p < 0.01$, **** $p < 0.0001$. All black dots represent individual animals.

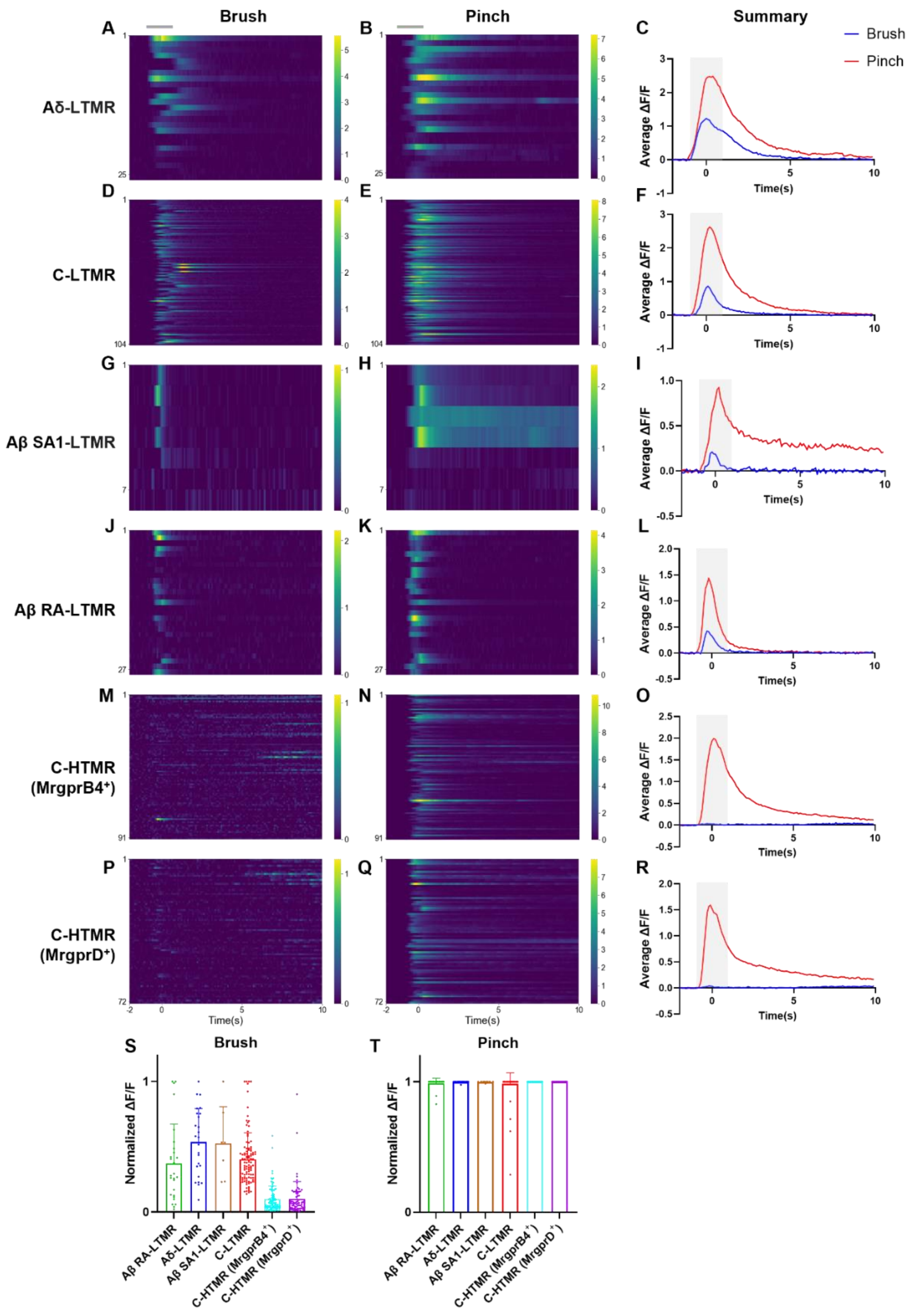


**Fig. S2. Brush and pinch responses of LTMRs and HTMRs**

(A-C) Heatmaps of calcium responses to (A) brush and (B) pinch stimuli, as well as (C) summary of averaged responses of Aδ-LTMR neurons.

(D-F) Heatmaps of calcium responses to (D) brush and (E) pinch stimuli, as well as (F) averaged responses of C-LTMR neurons.
(G-I) Heatmaps of calcium responses to (G) brush and (H) pinch stimuli, as well as (I) averaged responses of Aβ SA1-LTMR neurons.
(J-L) Heatmaps of calcium responses to (J) brush and (K) pinch stimuli, as well as (L) averaged responses of Aβ RA-LTMR neurons.
(M-O) Heatmaps of calcium responses to (M) brush and (N) pinch stimuli, as well as (O) averaged responses of C-HTMR (MrgprB4$^+$) neurons.
(P-R) Heatmaps of calcium responses to (P) brush and (Q) pinch stimuli, as well as (R) averaged responses of C-HTMR (MrgprD$^+$) neurons.
(S-T) Summaries of peak calcium responses to (S) brush and (T) pinch stimuli, normalized to each neuron's maximum responses to all stimuli. Dots represent individual neurons. Shown are the means ± SD. Same neurons and animals as in Fig. 2. The arrangement of neurons in rows in the heatmaps corresponds to the same arrangement shown in Fig. 2, with each row representing the same neuron across both figures.

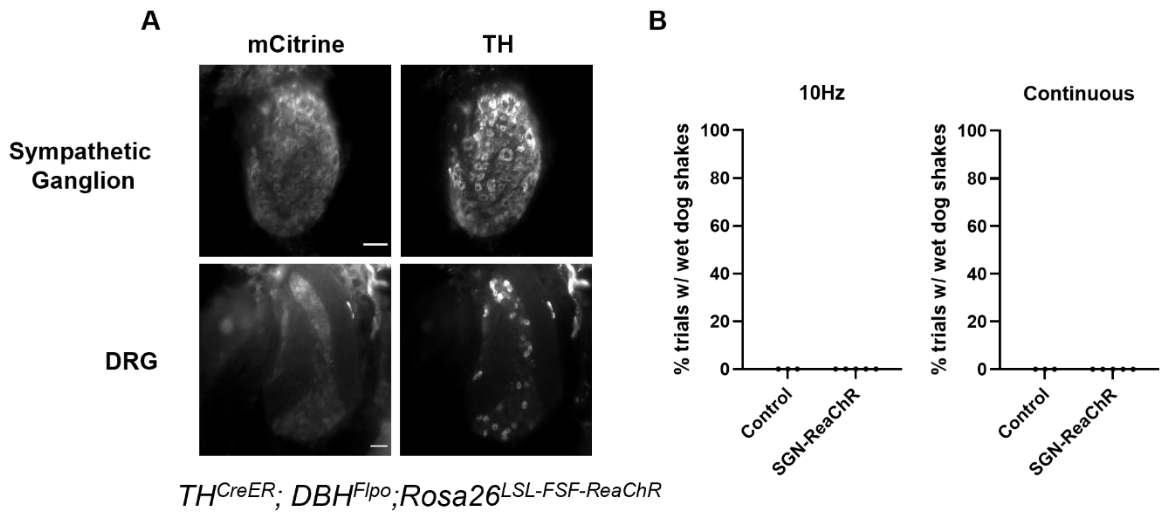


**Fig. S3. Optogenetic stimulation of sympathetic neurons terminating in the skin does not evoke wet dog shakes**

(A) Representative immunostaining images showing selective ReaChR (labeled by mCitrine) expression in the sympathetic ganglion neurons (SGNs) but not in the DRG neurons of $TH^{CreER}$; $DBH^{Flpo}$; $Rosa26^{LSL\text{-}FSF\text{-}ReaChR}$ animals. Scale bars, 50 μm.

(B) Percentage of successful trials with wet dog shakes induced by optogenetic stimulation of sympathetic neurons ($TH^{CreER}$; $DBH^{Flpo}$; $Rosa26^{LSL\text{-}FSF\text{-}ReaChR}$, N = 5) or control (littermates without CreER or Flpo allele, N = 3) on the neck. All markers represent individual animals.

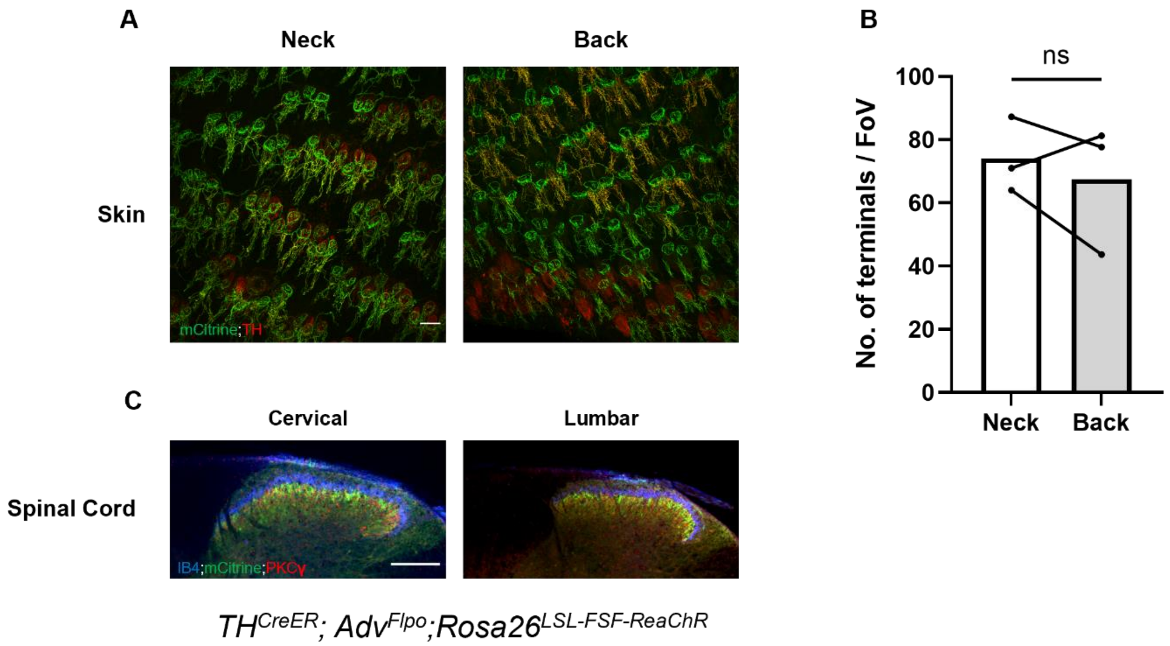


**Fig. S4. Comparison of C-LTMR terminals innervating the neck and lower back skin**

(A) Representative immunostaining images of C-LTMR lanceolate endings (labeled by mCitrine) that innervate the neck and the lower back hairy skin. Scale bars, 200 µm.
(B) Quantification of the average number of C-LTMR terminals per field of view (FoV) as represented in (A) across animals (N = 3). At least 3 images from different FoVs were averaged for each animal. Comparison was done using two-sided paired t test.
(C) Representative immunostaining images of C-LTMR central projections (labeled by mCitrine) that terminate in lamina IIi of the cervical and lumbar spinal cord. Scale bars, 200 µm. All markers represent individual animals. ns $p > 0.05$.

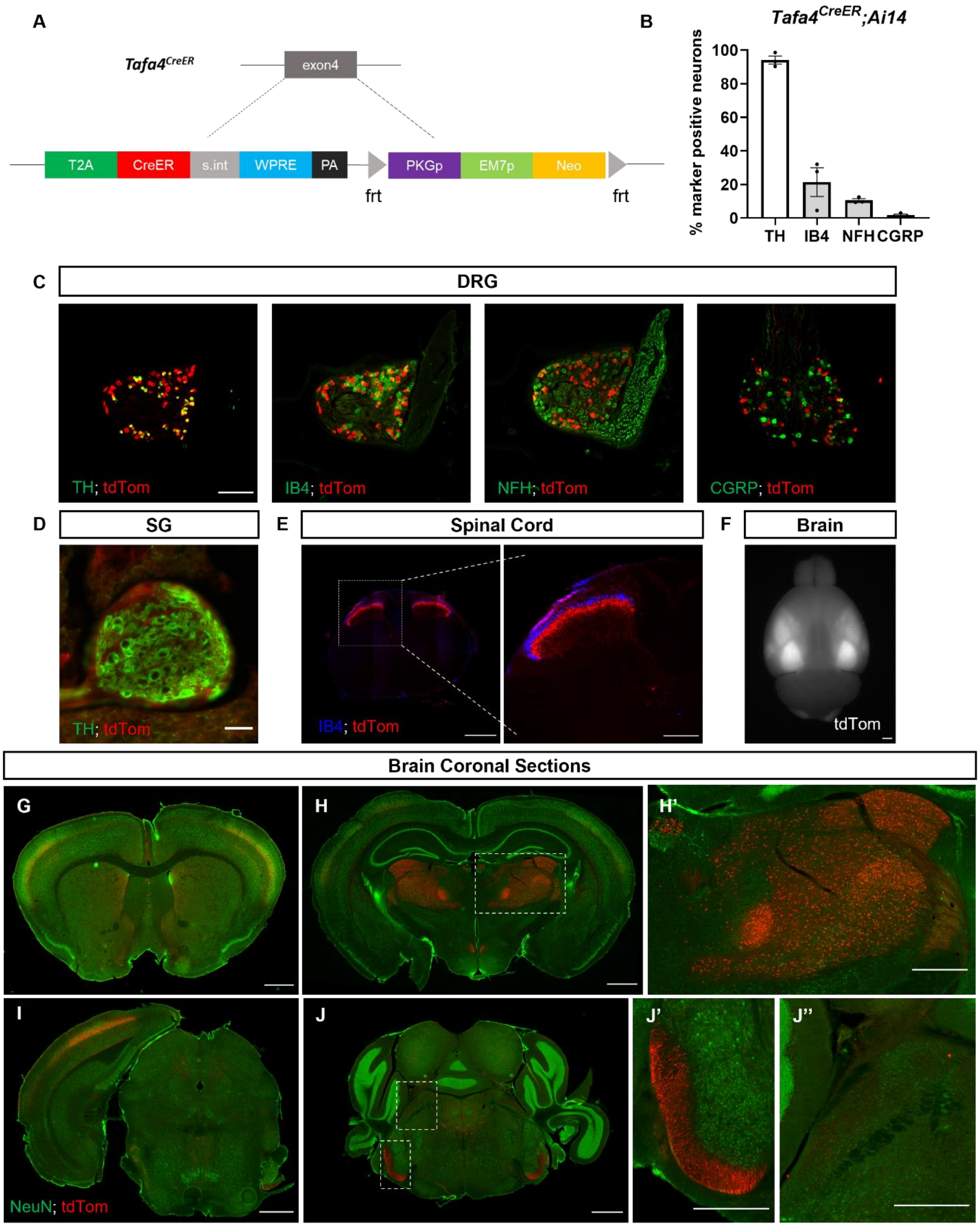


**Fig. S5. Generation and characterization of *Tafa4*$^{CreER}$ mice**

(A) Gene targeting strategy used to generate the *Tafa4$^{reER}$* mouse line. Exon annotation based on NCBI Reference Sequence accession NM_177233.5.
(B) Percentage of cervical DRG neurons labeled with specific cell type markers that are tdTomato positive in *TAFA4$^{CreER}$*; Ai14 animals (N =3). Dots represent individual animals, at least 3 sections were averaged from each animal. Shown are the means ± SEM.
(C) Representative images of cervical DRG neurons co-stained with cell type markers (green) and tdTomato (red). Scale bar, 200 μm.
(D) Representative image of sympathetic ganglion (SG) neurons co-stained with TH (green) and tdTomato (red). Scale bar, 50 μm.
(E) Representative images of cervical spinal cord sections co-stained with IB4 (blue) and tdTomato (red). Scale bar on the left panel, 500 μm. Right panel, a zoomed-in image of the dotted square region from the left panel. Scale bar, 200 μm.
(F) Representative images of whole brain toTomato florescence of a *Tafa4$^{CreER}$*; Ai14 animal. Scale bar: 1000 μm.
(G-J) Representative brain coronal sections containing (G) basal ganglion, (H) sensory thalamus including dLGN, PO, VPM, VPL, (I) midbrain, and (J) brainstem from *Tafa4$^{CreER}$*; Ai14 animals (scale bar: 1000 μm). Labeled cells are primarily seen in the thalamus across the entire brain. (H') is a zoomed-in image from (H) showing cell bodies of labeled neurons (scale bar, 500 μm) in the thalamus. (J', J'') are zoomed-in images from (J), showing (J') labeling of spinal tract of the trigeminal nerve, and (J'') no labeling of neurons in the parabrachial nucleus.

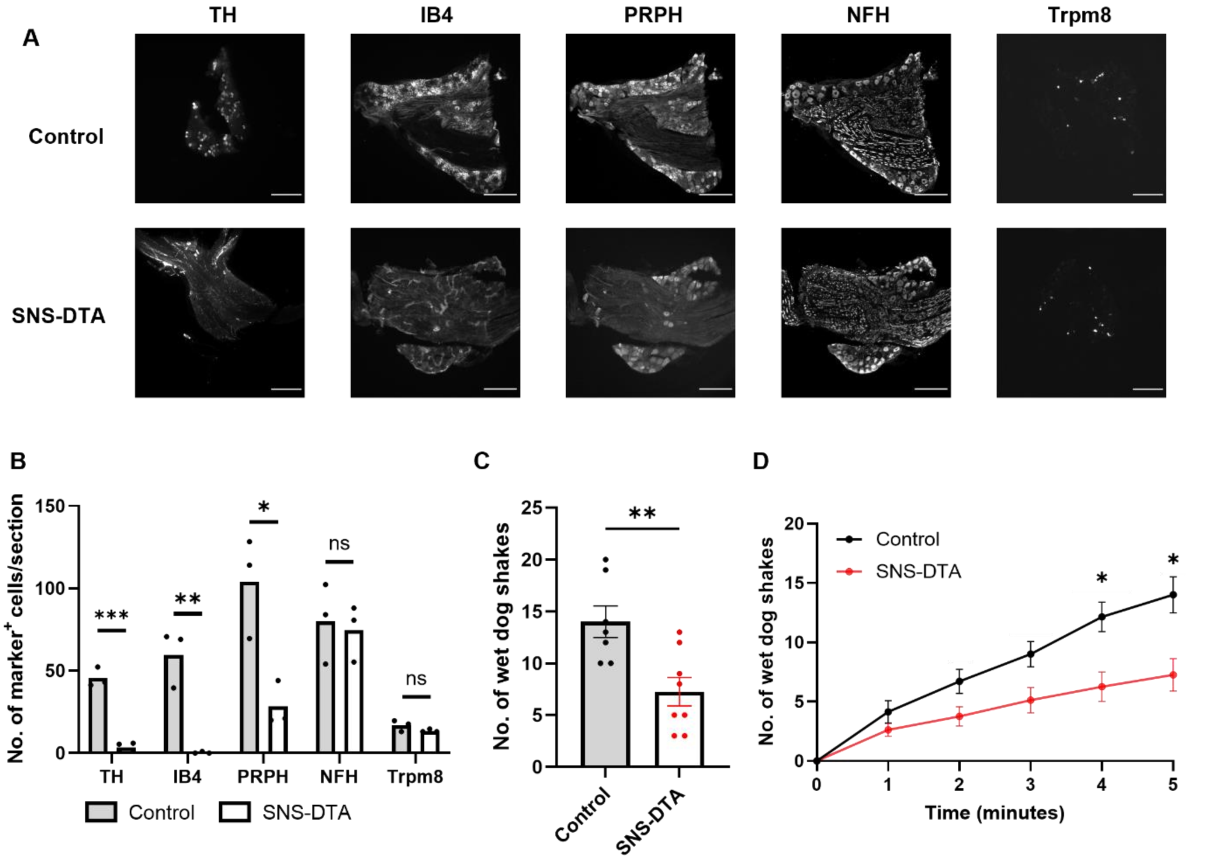


**Fig. S6. Oil droplet-induced wet dog shakes are reduced in SNS-DTA mice**

(A) Representative images of cervical DRG immunostaining of control (*Rosa26$^{LSL-DTA}$*) and SNS-DTA (*Scn10a$^{Cre}$; Rosa26$^{LSL-DTA}$*) animals. Scale bars, 200 μm.
(B) Average number of DRG neurons stained with specific cell type markers from (A). Each dot represents the average number of cells per section from one animal, at least 5 sections were averaged for each animal. Bars represent means. Comparison using two-sided unpaired t-test.
(C) Total number of oil-induced wet dog shakes in littermate control (N = 7) and SNS-DTA animals (N = 8) over 5 minutes. Dots represent individual animals. Comparison using two-sided unpaired t-test.
(D) Cumulative plot of the average number of wet dog shakes across animals over time. Same wet dog shake bouts as in (C). Comparisons were done using two-way ANOVA. p-values adjusted for multiple testing using Bonferroni correction. Shown are the means ± SEM. ns $p > 0.05$, * $p \leqslant 0.05$, ** $p < 0.01$, *** $p < 0.001$.

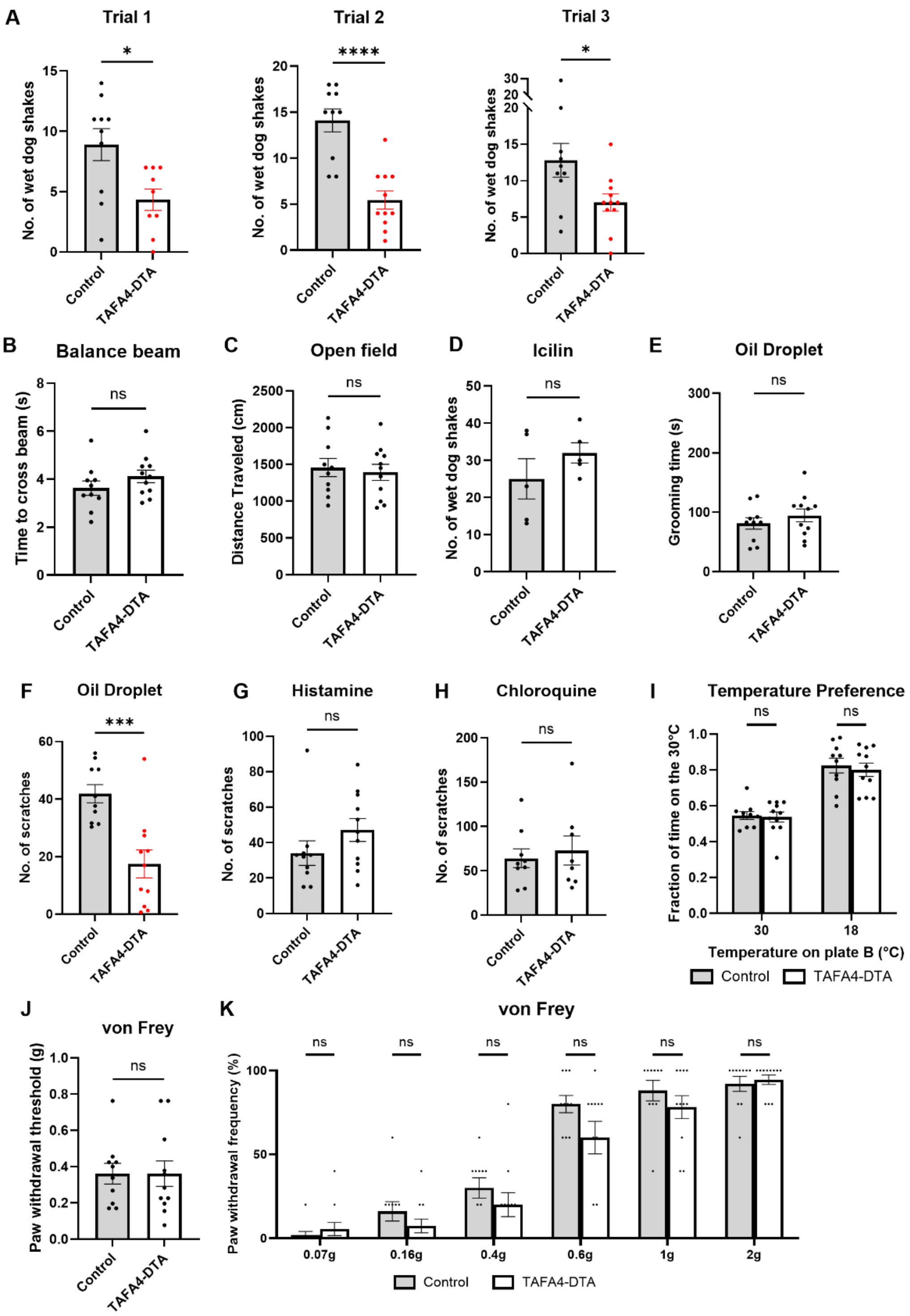


**Fig. S7. Additional behavioral responses of TAFA4-DTA mice**

(A) Total number of oil-induced wet dog shakes in littermate control (*Rosa26*$^{LSL\text{-}DTA}$, N = 10, 10, 10) and TAFA4-DTA animals over 5 minutes (*Tafa4*$^{CreER}$; *Rosa26*$^{LSL\text{-}DTA}$, N = 9, 11, 11) across 3 trials.

(B) Total time to cross the balance beam.
(C) Distance traveled in the open field test.
(D) Total number of icilin-induced wet dog shakes within 15 minutes post icilin administration. Control, N = 5; TAFA4-DTA, N = 5.
(E) Average grooming amount within 5 minutes (300 seconds) post oil droplet application. Time averaged across 3 trials in (A).
(F) Total number of scratching events within 5 minutes post oil droplet application averaged across 3 trials in (A).
(G) Total number of scratches within 15 minutes post histamine injection. Control, N = 5; TAFA4-DTA, N = 5.
(H) Total number of scratching events within 15 minutes post chloroquine injection. Control, N = 7; TAFA4-DTA, N = 5.
(I) Average fraction of time spent at 30 °C side of the chamber over a 5-minute period.
(J) Paw withdrawal threshold measured by 'up-down' von Frey method across animals.
(K) Percentage frequency of paw withdrawal (5 trials) in response to von Frey filament at different forces across animals. The same cohort of animals was used throughout all experiments. Number of control animals = 10, Number of TAFA4-DTA animals = 11, unless stated otherwise. Shown are the means ± SEM. All dots represent individual animals. Comparison was done using two-sided unpaired t-test. ns $p > 0.05$, * $p \leq 0.05$, ***$p < 0.001$, ****$p < 0.0001$

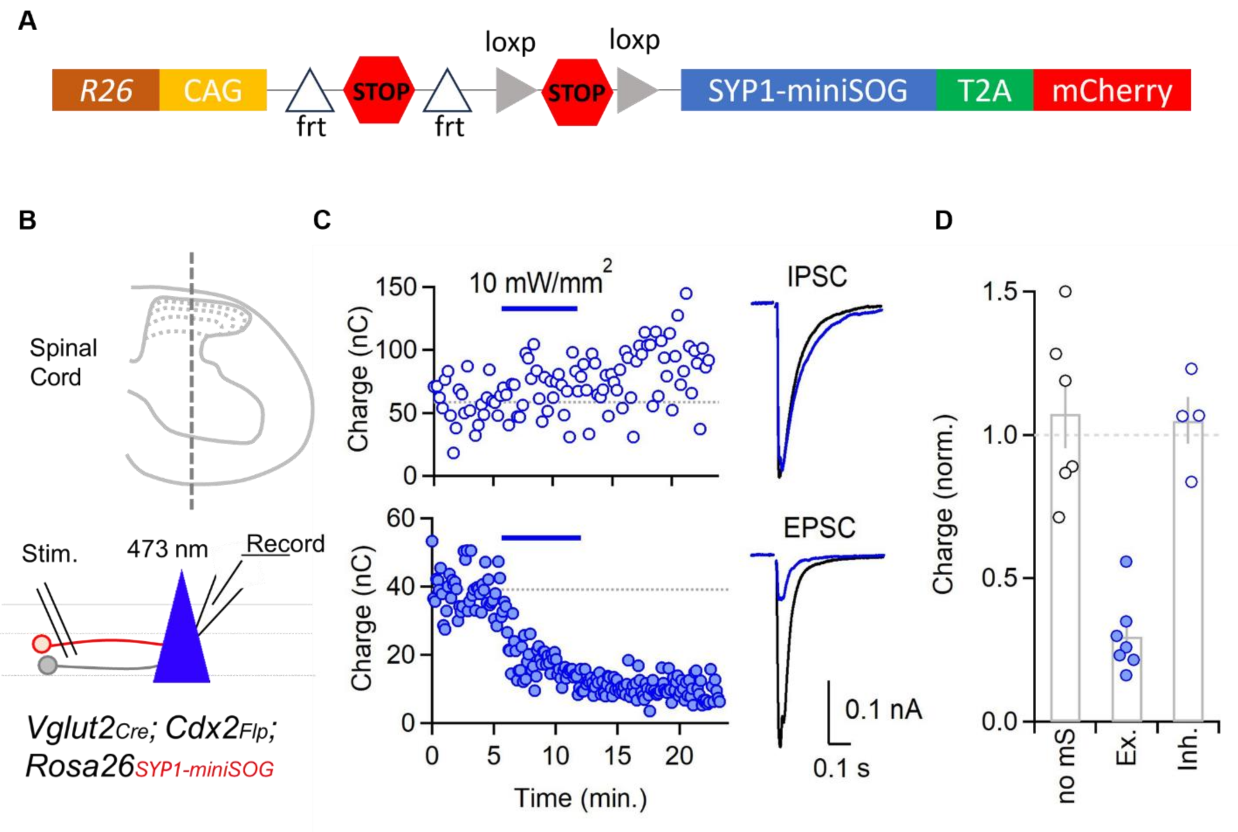


**Fig. S8. Generation and characterization of *Rosa26*$^{LSL\text{-}FSF\text{-}SYP1\text{-}mini\text{-}SOG\text{-}mcherry}$ mice**

(A) Gene targeting strategy used to generate the *Rosa26*$^{LSL\text{-}FSF\text{-}SYP1\text{-}mini\text{-}SOG\text{-}mcherry}$ mouse line.
(B) Schematic diagram showing genetic manipulation strategies and experimental setup. Random lamina II-IV spinal dorsal horn neurons were recorded while stimulating neighboring cells in a sagittal spinal cord slice. Blue light was delivered to the cell body using a 40x objective.
(C) Example cells in which IPSCs (top left) and EPSCs (bottom left) were recorded. Light was then delivered to suppress excitatory synapses. Each dot indicates a trial. Averaged evoked currents before (black) and after light stimulation (blue) for these two cells are shown at right.
(D) Normalized synaptic charge of control (animals without miniSOG expression, no mS) and miniSOG expressing animals after light stimulation. Synaptic charges are normalized to evoked charge before light stimulation. All dots represent individual cells. Bars represent means.

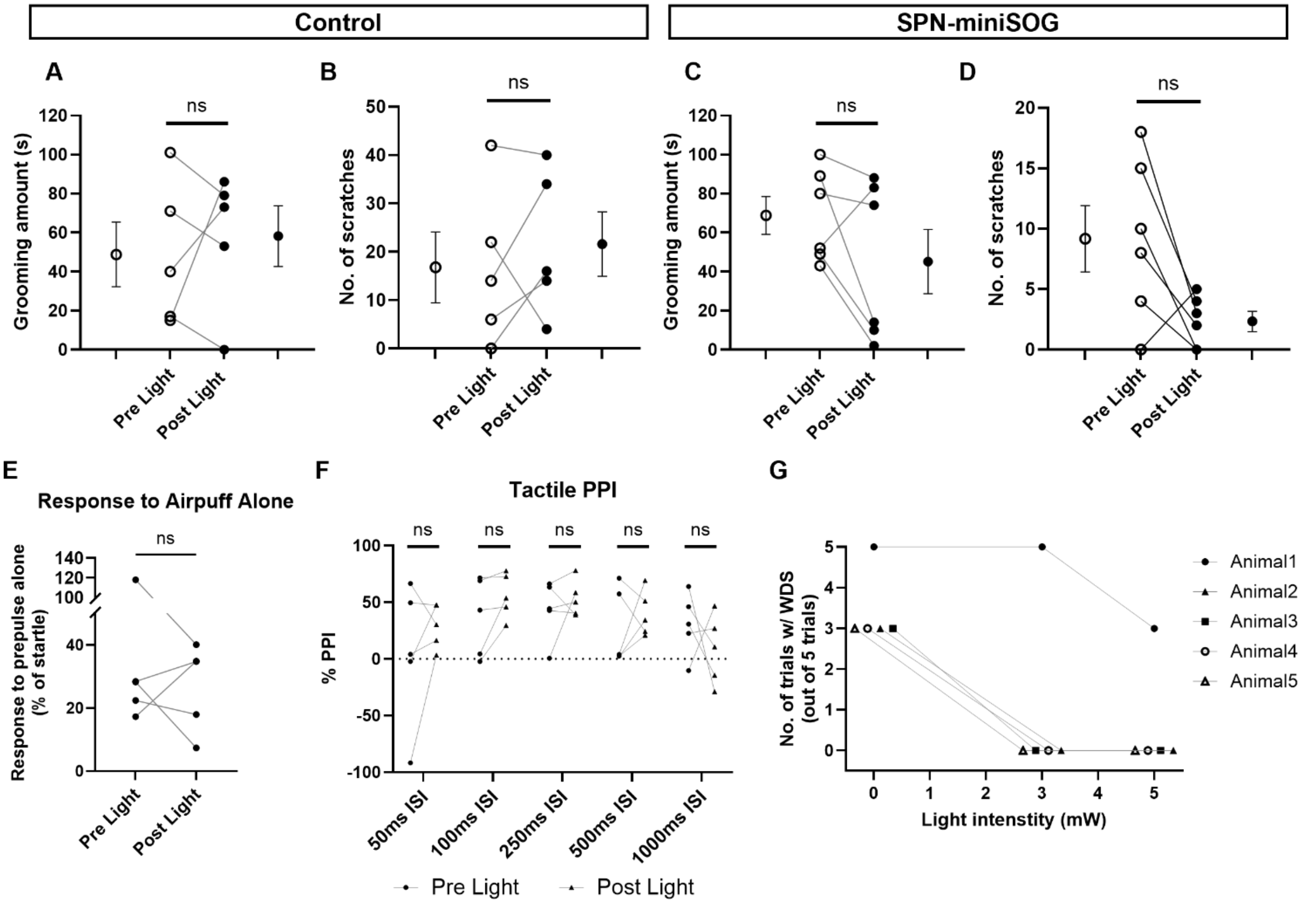


**Fig. S9. Light effects on oil droplet-induced behaviors in Vgat-ChR2 and SPN-miniSOG animals**

(A-D) Quantification of (A, C) the grooming amount and (B, D) the number of scratching events within 5 minutes post oil droplet application in littermate control (N = 5) and SPN-miniSOG animals (N = 6). Dots in the middle represent individual animals. Dots on the lateral sides represent the average responses. Shown are the means ± SEM. ns $p > 0.05$
(E) Response to a light air puff stimulus alone (0.9 PSI, 50 ms) before and after light stimulation in SPN-miniSOG animals.
(F) Percent inhibition of the startle response at multiple interstimulus intervals (ISIs) between the prepulse (0.9 PSI air puff) and the pulse (125-dB sound) in SPN-miniSOG animals before and after light stimulation.
(G) Relationship between light intensity (brain) and the number of trials with evoked wet dog shakes during $TH^+$ skin afferent photoactivation across animals. Light intensities tested include 0 mW (no light), 3 mW, and 5 mW.

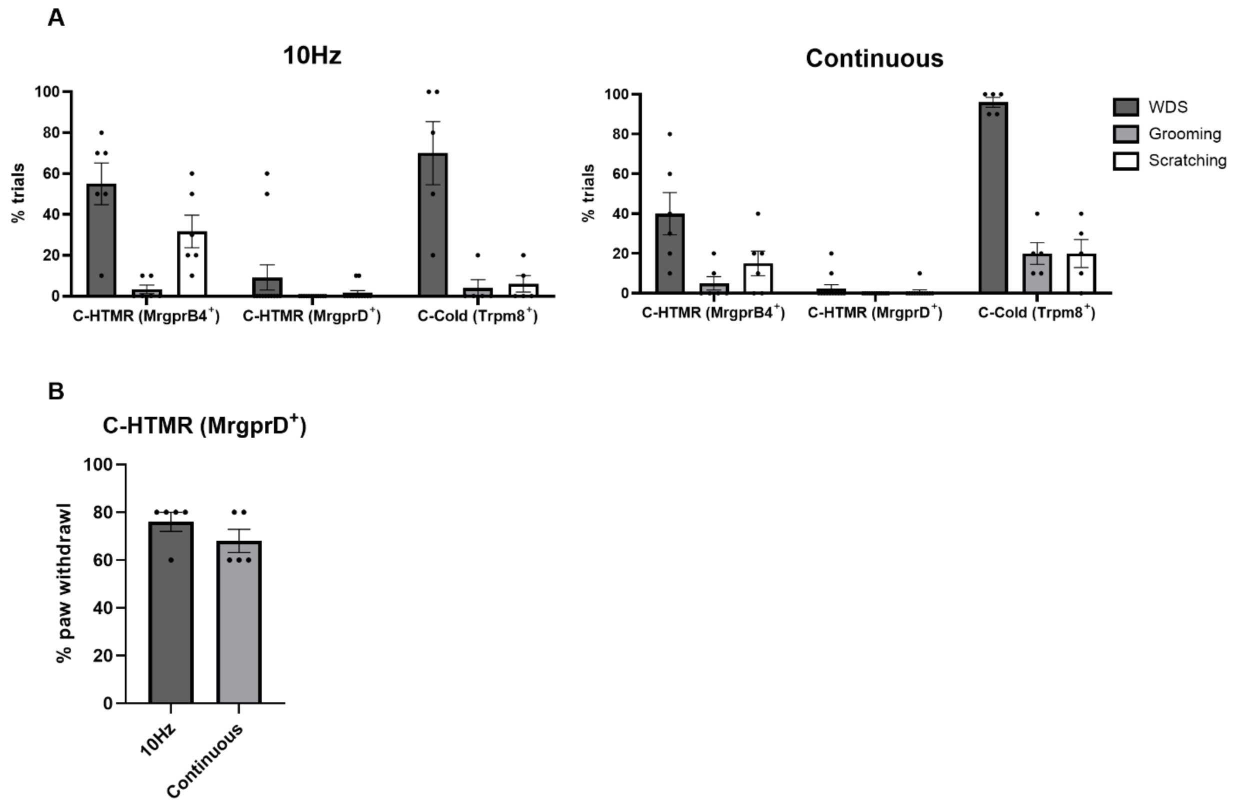


**Fig. S10. Optogenetic stimulation of C-HTMRs (MrgprB4$^{+}$), C-HTMRs (MrgprD$^{+}$), and C-Cold (Trpm8$^{+}$) neurons**

(A) Percentage of successful trials with wet dog shakes induced by illuminating light on the neck skin of *MrgprB4$^{Cre}$; Rosa26$^{LSL\text{-}ReaChR}$* (N = 6), *MrgprD$^{CreER}$; Rosa26$^{LSL\text{-}ReaChR}$* (N = 12), and *Trpm8$^{Flpo}$; Rosa26$^{FSF\text{-}ReaChR}$* (N = 5) animals.

(B) Percentage of trials with light-induced paw withdrawal responses. Data collected from a subset of *MrgprD$^{CreER}$; Rosa26$^{LSL\text{-}ReaChR}$* animals in (A) (N = 5) that did not show light induced wet dog shakes, indicating functional expression of ReaChR in MrgprD$^{+}$ neurons. A total of 5 trials were conducted for each type of light stimulation paradigm. All markers represent individual animals. Shown are the means ± SEM.
